# A genetically encoded microtubule bundler for causal dissection of microtubule bundling in cells

**DOI:** 10.64898/2026.01.22.700042

**Authors:** Soei Watari, Takumi Chinen, Yuto Kunitatsu, Takashi Saitou, Yasufumi Takahashi, Kazunori Matsuura, Takanari Inoue, Hiroshi Inaba

**Affiliations:** Department of Chemistry and Biotechnology, Graduate School of Engineering, Tottori University, Tottori 680-8552, Japan; Department of Cell Biology and Center for Cell Dynamics, Johns Hopkins University School of Medicine, 855 North Wolfe Street, Baltimore, MD 21205, USA; Graduate School of Pharmaceutical Sciences, The University of Tokyo, 7-3-1 Hongo, Bunkyo-ku, Tokyo 113-8654, Japan; Department of Electrical Engineering, Graduate School of Engineering, Nagoya University, Aichi 464-8601, Japan; Graduate School of Science and Engineering, Ehime University, 3 Bunkyo-cho, Matsuyama, Ehime 790-8577, Japan; Research Institute for Quantum and Chemical Innovation, Institutes of Innovation for Future Society, Nagoya University, Nagoya 464-8601, Japan; WPI Nano Life Science Institute (WPI-NanoLSI), Kanazawa University, Kanazawa 920-1192, Japan; Centre for Research on Green Sustainable Chemistry, Tottori University, Tottori 680-8552, Japan; Chromosome Engineering Research Center, Tottori University, Tottori 683-8503, Japan; Graduate School of Integrated Sciences for Life, Hiroshima University, 1-3-1 Kagamiyama, Higashi-Hiroshima, Hiroshima 739-8526, Japan

## Abstract

Microtubule (MT) bundling is a conserved organizational feature of the cytoskeleton that accompanies MT stabilization, suggesting that it is involved in diverse cellular processes, including mitosis, migration, and axon morphogenesis. Although microtubule-associated proteins are known to induce MT bundling, whether bundling itself is sufficient to alter MT properties and cellular behavior has remained difficult to address due to the lack of tools that selectively manipulate MT bundling in living cells. Here, we describe the development of a genetically encoded, protein-based “MT-Bundler” by coupling an MT-binding motif to a biologically inert oligomerization scaffold, enabling direct and tunable crosslinking of intracellular MTs. The expression of MT-Bundler consisting of MAP4 and Azami-Green, drove robust MT bundling, but also conferred marked resistance to depolymerization, and elevated MT acetylation. Functionally, enforced MT bundling disrupts cell division and migration and suppresses neurite and axon outgrowth. To confirm the causal relationship behind these findings, we further engineered chemically and optically inducible MT-Bundlers that permit rapid, reversible, and spatiotemporally precise control of MT bundling. Acute induction of MT bundling triggers a rapid increase in MT acetylation, revealing bundling as an upstream organizational cue that promotes luminal access of the acetyltransferase ATAT1. Notably, MT stabilization persists even in the absence of acetylation, demonstrating that bundling itself is sufficient to mechanically stabilize MTs. Together, these results identify MT bundling as a primary determinant of MT stability and modification, establishing MT-Bundlers as a versatile tool to dissect the mechanistic basis of MT bundling in living cells.

## INTRODUCTION

Microtubules (MTs) are dynamic cytoskeletal polymers that organize intracellular architecture and enable essential cellular processes, including cell division, migration, and neuronal morphogenesis^1–5^. Beyond their intrinsic dynamics, MTs are assembled into higher-order structures—such as bundles—that are characteristic of specialized cellular states^6,7^. Prominent examples include parallel MT arrays in axons and dendrites, bundled MTs in the spindle midzone, and stabilized MT networks in the perinuclear region^8^. MT bundling is widely associated with enhanced MT stability and long-lived cellular structures; however, whether bundling itself directly alters MT properties or instead reflects secondary consequences of microtubule-associated proteins (MAPs) activity remains unresolved.

In cells, MT bundling is mediated by diverse MAPs that differ in structure, localization, and regulatory roles^9,10^. In neurons, MT crosslinking by MAP2 and Tau organizes parallel MT arrays that support axon and dendrite integrity and growth^11,12^. In non-neuronal cells, MAP4 promotes MT bundling preferentially at the cell periphery, contributing to cell shape regulation and protrusion formation^13–15^. Additional MT crosslinking factors, including PRC1, play essential roles during mitosis by organizing bundled MTs in the spindle midzone^16^. More recently, members of the microtubule crosslinking factor family, MTCL1 and MTCL2, have been shown to stabilize bundled MTs in the perinuclear region and at the Golgi apparatus, contributing to asymmetric MT organization and intracellular trafficking^8,17,18^. Although these MAPs clearly demonstrate the physiological importance of MT bundling, they typically perform multiple functions simultaneously, including MT binding, crosslinking, regulation of dynamics, and recruitment of additional factors. As a result, it has been difficult to disentangle the direct effects of MT bundling itself from other MAP-dependent activities.

Pharmacological approaches have provided powerful tools to perturb MT polymerization and depolymerization and have been instrumental in elucidating MT-dependent processes such as spindle checkpoint signaling and organelle positioning^19–22^. However, most MT-targeting drugs do not recapitulate the large-scale architectural rearrangements associated with MT bundling, limiting their utility for probing the structural consequences of bundled MT organization. Consequently, the direct relationship between MT bundling, MT stability, and cellular function remains incompletely understood.

One prominent correlate of bundled MTs is increased acetylation of α-tubulin at lysine 40, a luminal post-translational modification catalyzed by the acetyltransferase ATAT1^23,24^. MT acetylation has been linked to long-lived MTs and implicated in processes such as cell migration, intracellular transport, and neuronal function^25–27^. However, whether acetylation is a primary driver of MT stabilization or instead arises as a consequence of altered MT architecture remains debated. Recent studies suggest that lattice defects, curvature, or expansion may facilitate luminal enzyme access^23,28,29^, raising the possibility that higher-order MT organization itself could promote acetylation. Progress in addressing these issues has been limited by the lack of tools that selectively and reversibly induce MT bundling in living cells, independently of endogenous MAP regulation. While protein-based multimerization strategies have been used to crosslink MTs *in vitro*^30,31^, a genetically encoded and temporally controllable MT-bundling tool suitable for live-cell studies has been lacking.

Here, we introduce a genetically encoded “MT-Bundler,” engineered by coupling a minimal MT-binding motif to a biologically inert oligomerization scaffold. This design enables direct crosslinking of intracellular MTs without perturbing endogenous MAP expression or enzymatic pathways. Using constitutive, chemically inducible, and optogenetic variants of the MT-Bundler, we systematically dissect the structural and functional consequences of MT bundling in living cells. We show that MT bundling is sufficient to stabilize MTs independently of acetylation, while also acting as an upstream structural trigger for rapid MT acetylation. Together, these findings establish MT bundling as a primary determinant of MT stability and modification and provide a versatile experimental framework for probing MT architecture and function *in vivo*.

## RESULTS AND DISCUSSION

### Design of MT-Bundler and analysis of its binding to MTs

To engineer a genetically encoded tool capable of inducing MT bundling, we employed the minimal MT-binding domain of MAP4 (MAP4m). MAP4m is a well-characterized motif, widely utilized as a reliable marker of MT organization when fused to monomeric fluorescent proteins^32,33^. To drive MT crosslinking, we selected the tetrameric fluorescent protein Azami-Green (AG) as a structural scaffold, reasoning that the multivalent display of MAP4m on this oligomeric core would facilitate bundle formation. We engineered the “MT-Bundler” by fusing MAP4m to the N-terminus of AG (MAP4m-AG) (Fig. 1a and Supplementary Fig. 1). Upon transient expression in HeLa cells, confocal microscopy revealed that MAP4m-AG strictly colocalized with the MT marker Tubulin Tracker Deep Red, inducing striking reorganization of the MT network into a bundled architecture (Fig. 1c). Line scan analysis confirmed a significant increase in tubulin fluorescence intensity along these filaments in MAP4m-AG-expressing cells, providing quantitative support for the formation of higher-order bundles (Fig. 1d). This bundling capability was shown not to be affected by topology, as fusion of MAP4m to the C-terminus of AG (AG-MAP4m) induced a similar phenotype (Supplementary Fig. 1 and 2). Furthermore, morphometric analysis indicated a reduction in the MT-occupied area relative to the total cell area (Fig. 1e), consistent with the spatial compaction of MTs via bundling. To clarify the fine structure of these induced assemblies at high resolution, we employed super-resolution imaging using stochastic optical reconstruction microscopy (STORM). Immunostaining of α-tubulin in MAP4m-AG-expressing cells revealed well-defined MT structures exhibiting distinct branched arrangements (Fig. 1f, red arrows). In contrast, expression of the AG scaffold without MAP4m resulted in diffuse MT fluorescence and the absence of organized MT networks (Supplementary Fig. 3). Collectively, these results demonstrate that MAP4m-AG effectively induces the formation of MT bundles in living cells.

**Fig. 1.**
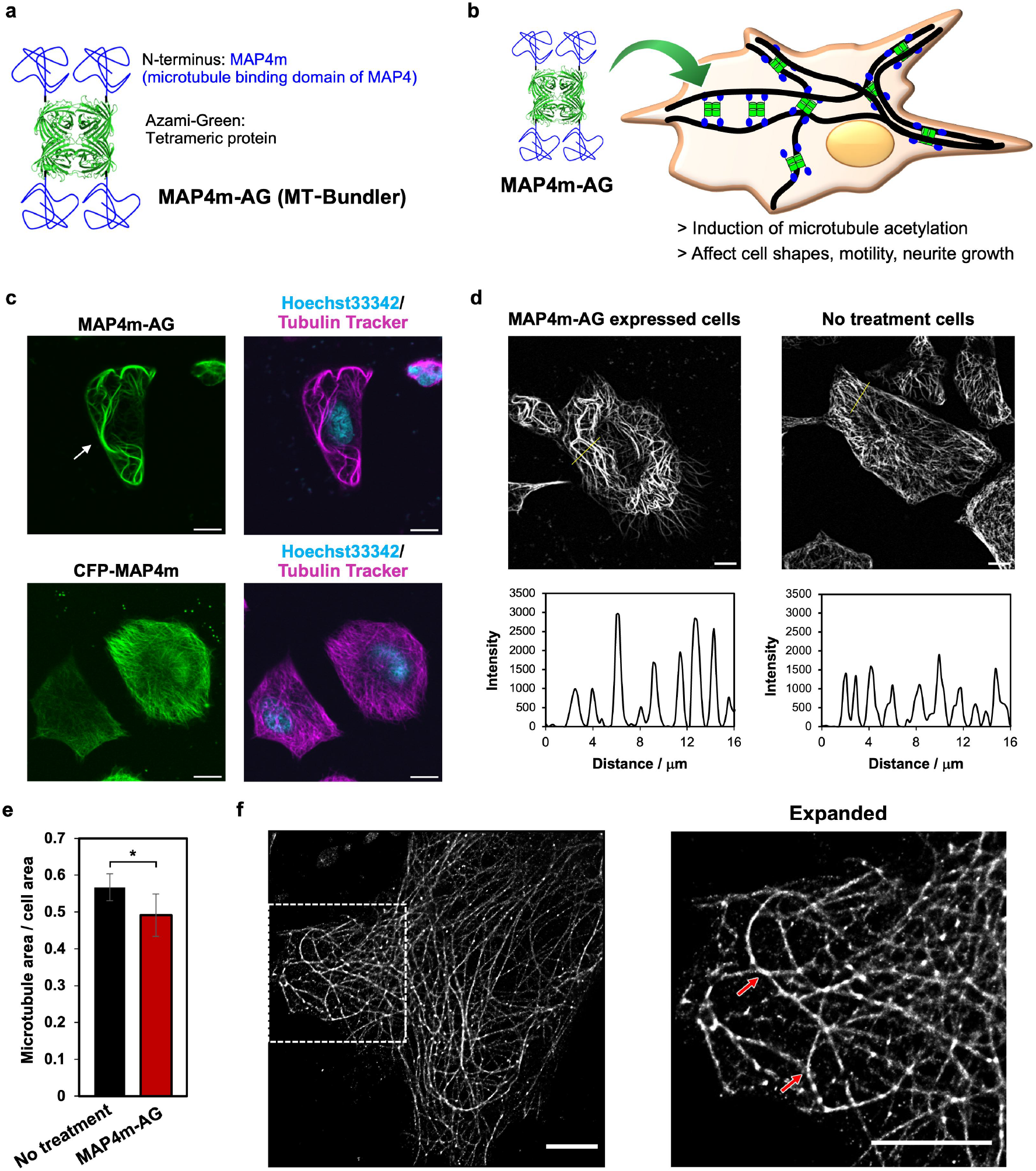
Design of microtubule (MT) bundler that crosslinks intracellular MTs. (a) Components of MT-Bundler, which is constructed with the MT binding domain of MAP4 (MAP4m) and the fluorescent tetrameric protein Azami-Green (AG). (b) Crosslinking of intracellular MTs by MT-Bundler in cells. (c) Confocal laser scanning microscopy images of MAP4m-AG-expressing HeLa cells (top). CFP-MAP4m was expressed as a control (bottom). MTs were stained with Tubulin Tracker Deep Red and cell nuclei were stained with Hoechst 33342. Scale bars, 10 μm. (d) Line profiling of fluorescence intensity of Tubulin Tracker Deep Red with or without MAP4m-AG expression. Scale bars, 10 μm. (e) Quantification of MT area per cell area with or without MAP4m-AG expression. Error bars represent the SD (*N* = 30). \**P* < 0.0001, two-tailed Student’s t-test. (f) Stochastic optical reconstruction microscopy (STORM) images of MAP4m-AG-expressing HeLa cells. The samples were stained with β-tubulin antibody. Red arrows indicate the branching of MTs. Scale bars, 5 μm.

### Tracking of MT bundling dynamics

To monitor the temporal dynamics of MT bundling, we performed live-cell time-lapse imaging using retinal pigment epithelium 1 (RPE1) cells stably expressing MAP4m-AG under a doxycycline-inducible Tet-On system. Imaging was initiated immediately upon doxycycline addition, allowing us to visualize the induction of MAP4m-AG expression and subsequent MT reorganization. MAP4m-AG bound to MTs immediately following its expression, rapidly triggering the formation of MT bundles (Fig. 2a and Supplementary Video 1). Notably, bundling was initiated preferentially in the perinuclear region, likely reflecting the higher density of MTs in this area, which facilitates crosslinking. As bundling progressed, cells exhibited pronounced elongation and morphological aberrations during cell division (Fig. 2b). Consistent with these structural defects, MAP4m-AG expression frequently resulted in mitotic arrest (Supplementary Video 1) and significantly suppressed cell proliferation as assessed by WST assays (Supplementary Fig. 4). These results indicate that excessive MT bundling interferes with essential mitotic machinery and cell cycle progression.

**Fig. 2.**
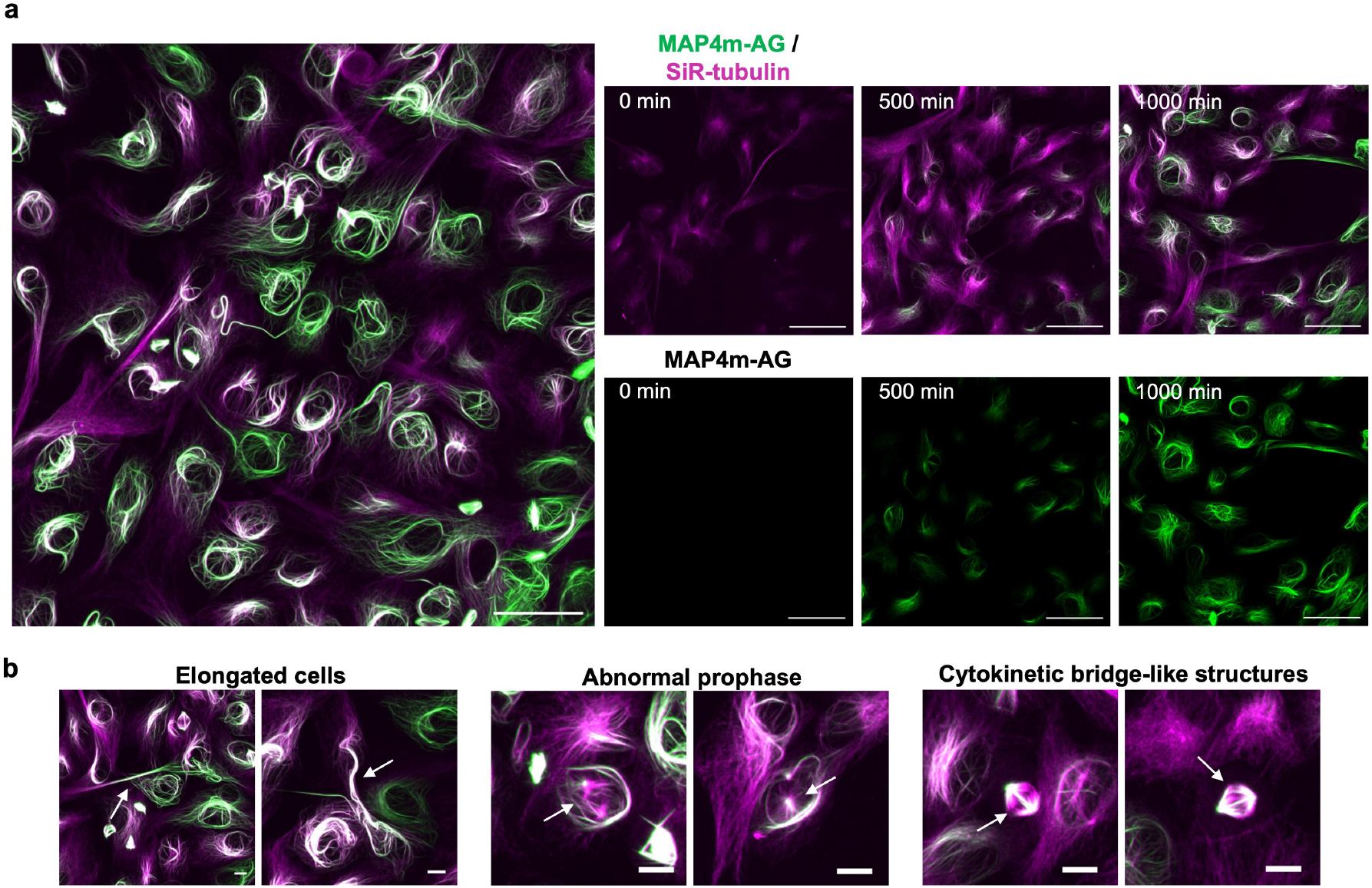
Trace of bundling of MTs by MT-Bundler and effects on cell structures. (a) Time-lapse fluorescence images show MAP4m-AG expression after 23 h of treatment with 1 μg/mL doxycycline. MTs were stained with SiR-tubulin. Scale bars, 50 μm. See also Supplementary Video 1. (b) Fluorescence images of abnormal phenotypes involving elongated cell shapes, abnormal prophase and cytokinetic bridge-like structures (white arrows). Scale bars, 10 μm.

### Effect of MT bundling on stability and acetylation of MTs

MT bundling is often associated with enhanced stability, although it remains unclear whether this arises from physical reinforcement or as a secondary consequence of acetylation. To disentangle these factors, we first evaluated structural stability by treating HeLa cells expressing MAP4m-AG with the MT depolymerizing agent nocodazole. While MTs in control cells expressing AG alone or monomeric MAP4m-fused monomeric AG (MAP4m-mAG) were largely depolymerized, bundled MTs in MAP4m-AG-expressing cells maintained their structure (Fig. 3a and Supplementary Fig. 5). Similar resistance to depolymerization was observed under cold treatment (4 °C), a condition known to destabilize dynamic MTs (Supplementary Fig. 6). These observations confirm that MAP4m-AG-mediated crosslinking confers significant structural stability to MTs.

**Fig. 3.**
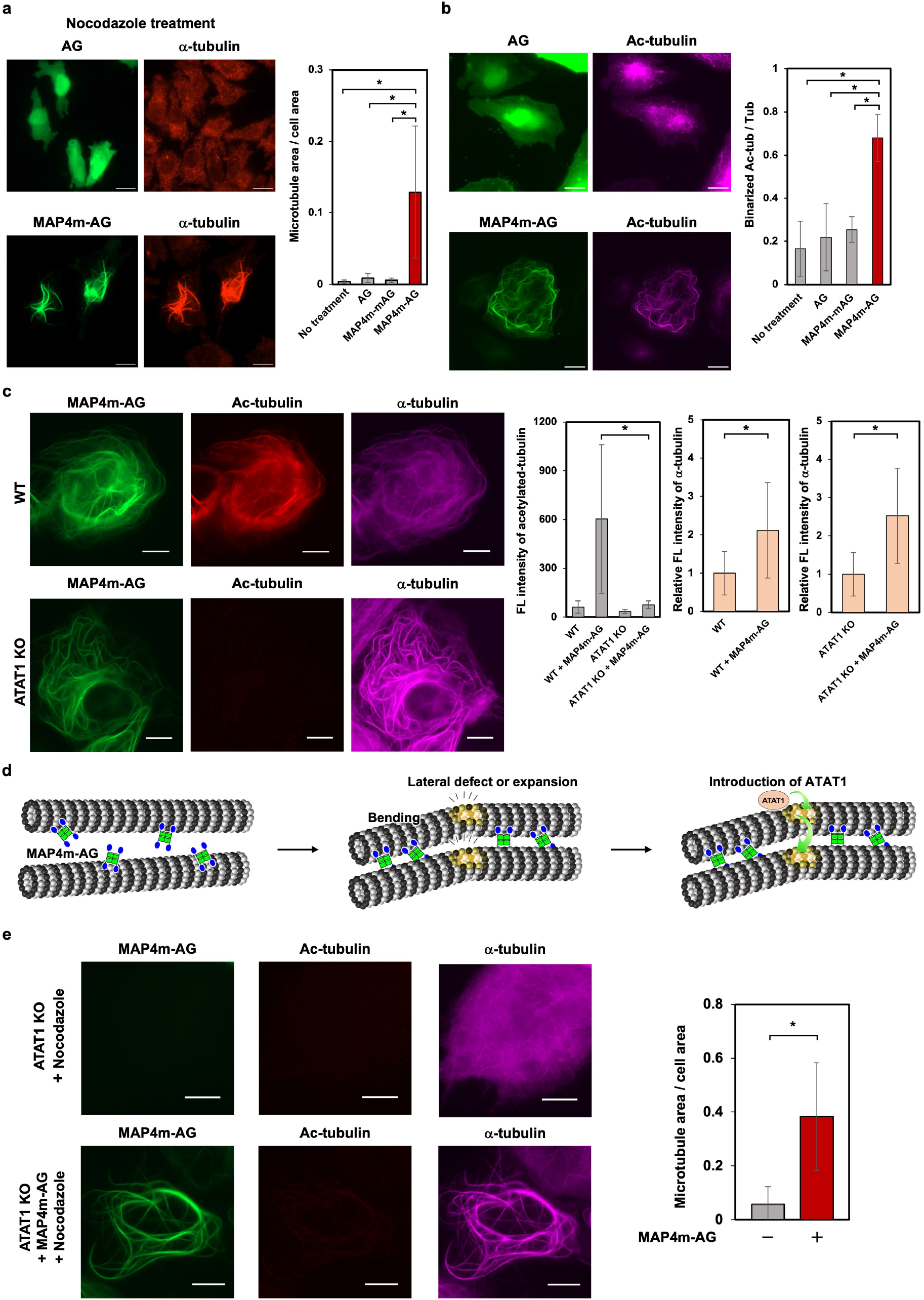
MAP4m-AG stabilizes MT structures and enhances acetylation level of MTs. (a) Fluorescence images of AG- or MAP4m-AG-expressing cells after treatment with 1 μM nocodazole for 2 h. HeLa cells transfected with AG or MAP4m-AG (green) were immunostained for α-tubulin (red). Scale bars, 20 μm. The graph shows the values of MT area per cell area. Error bars represent the SD (*N* = 13). \**P* < 0.0001, two-tailed Student’s t-test. (b) Acetylation level of MTs in AG or MAP4m-AG-expressing cells. HeLa cells transfected with tubulin-mCherry and AG or MAP4m-AG (green) were immunostained for acetylated (Ac)-tubulin (magenta). Scale bars, 20 μm. The graph shows the ratio of Ac-tubulin per mCherry-labeled MT. Error bars represent the SD (*N* = 5). \**P* < 0.001, two-tailed Student’s t-test. (c) Fluorescence images of MAP4m-AG-expressing RPE1 cells with or without ATAT1 knockout. Ac-tubulin and α-tubulin were immunostained. Scale bars, 10 μm. Graphs show the fluorescence intensity of Ac-tubulin (left) and, the relative fluorescence intensity of α-tubulin (middle: WT, right: ATAT1 KO). Error bars represent the SD (*N* = 20). \**P* < 0.001, two-tailed Student’s t-test. (d) Estimated mechanism of bunding and subsequent acetylation of MTs. (e) Fluorescence images of nocodazole-treated RPE1 cells with or without MAP4m-AG expression with ATAT1 knockout. Ac-tubulin and α-tubulin were immunostained. Scale bars, 10 μm. The graph shows the values of MT area per cell area. Error bars represent the SD (*N* = 11). \**P* < 0.0001, two-tailed Student’s t-test.

We next examined α-tubulin acetylation at lysine 40 (K40), a representative post-translational modification catalyzed by the acetyltransferase ATAT1 and associated with stable, long-lived MTs^25–27^. Immunofluorescence and western blot analyses revealed the marked upregulation of acetylated (Ac)-tubulin levels in MAP4m-AG-expressing cells compared with the level in controls (Fig. 3b, Supplementary Fig. 7, and Supplementary Fig. 8). To determine the hierarchy between bundling and acetylation, we employed ATAT1-knockout RPE1 cells. To this end, ATAT1 was deleted in RPE1 cells using a CRISPR-del strategy that removes the entire gene locus (Supplementary Fig. 9). In the absence of ATAT1, MAP4m-AG still induced robust MT bundling, despite the complete loss of acetylation signals (Fig. 3c). Quantitative analysis corroborated that, while acetylation was abolished in the knockout cells, the extent of bundling (measured by tubulin intensity) remained unaffected. This demonstrates that MT bundling occurs independently of acetylation. To assess whether the artificial MT bundling induced by MAP4m-AG mimics the physiological properties of native MAPs, we employed mAG-Tau382, a mAG-fused isoform of the canonical MT-bundling protein Tau (Supplementary Fig. 1). Consistent with the observation using MAP4m-AG, transient expression of mAG-Tau382 induced prominent MT bundling in RPE1 cells (Supplementary Fig. 10). Notably, this bundling was unaltered in ATAT1-knockout cells, demonstrating that the artificial bundles induced by MAP4m-AG mimic the acetylation-independent behavior of naturally occurring MT bundles. Based on these findings, we propose a model in which bundling acts as the upstream structural signal (Fig. 3d). Specifically, MAP4m-AG-induced crosslinking likely generates lattice curvature, defects, or expansion, which in turn facilitates the luminal entry of ATAT1 to drive acetylation as a secondary consequence^23,28,29^. Because various MAPs, such as Tau, induce similar bundling, this structure-dependent accessibility mechanism may be a conserved feature of cytoskeletal regulation. Crucially, to determine whether bundling itself is sufficient for stability, we treated ATAT1-knockout cells expressing MAP4m-AG with nocodazole. Remarkably, bundled MTs retained their resistance to depolymerization, even in the absence of acetylation (Fig. 3e). This finding challenges the prevailing view that acetylation is a requirement for stability in these structures, suggesting instead that physical crosslinking by MAP4m-AG mechanically reinforces MTs to obstruct their disassembly.

### Functional consequences on cell morphology and migration

We next assessed the impact of enforced MT bundling on cellular architecture and physiology. To evaluate the alterations in the mechanical properties of cells induced by MT bundling, we employed scanning ion conductance microscopy (SICM)^34^. Analysis of COS-7 cells expressing MAP4m-AG revealed a significant increase in mechanical stiffness specifically within regions containing bundled microtubules (Fig. 4a). Furthermore, these cells exhibited pronounced spatial heterogeneity in their mechanical properties across different cellular regions. In contrast, control cells expressing AG or MAP4m-mAG did not display such localized increases in stiffness (Supplementary Fig. 11). In these control cells, the mechanical profile remained relatively uniform, indicating suppression of the mechanical heterogeneity observed in the MT-bundled cells. Single-cell tracking revealed that, while control cells (expressing AG or MAP4m-mAG) displayed normal motility, MAP4m-AG expression significantly reduced the velocity of cell migration (Fig. 4b, Supplementary Video 2). Given that MT disassembly is necessary for focal adhesion turnover^35^, these results suggest that excessive stabilization freezes the cytoskeletal dynamics necessary for efficient migration. HeLa cells expressing MAP4m-AG adopted a distinctive elongated morphology (Fig. 4c, Supplementary Fig. 12, Supplementary Video 3), a phenotype likely driven by the longitudinal alignment of stiffened MT bundles^36^. To evaluate the impact on neuronal morphogenesis, we examined neurite outgrowth in Neuro-2a cells, a process normally orchestrated by MAPs such as Tau^37^. Although cells expressing the MT plus-end tracker EB3-tdTomato (a positive control) exhibited robust neurite extension^38^, MAP4m-AG expression potently suppressed outgrowth (Fig. 4d). Quantitative analysis showed a dramatic shift in population distribution: The proportion of cells lacking neurites increased from 72% to 94%, while those with extended neurites dropped from 28% to 6.3% (Fig. 4e). This phenotype mirrors the effects of DDA3, a physiological suppressor of neurite formation known to act via MT bundling^38^. These findings suggest that artificially induced hyper-bundling disrupts the delicate balance of endogenous MAPs required for axonogenesis, effectively stalling neuronal development.

**Fig. 4.**
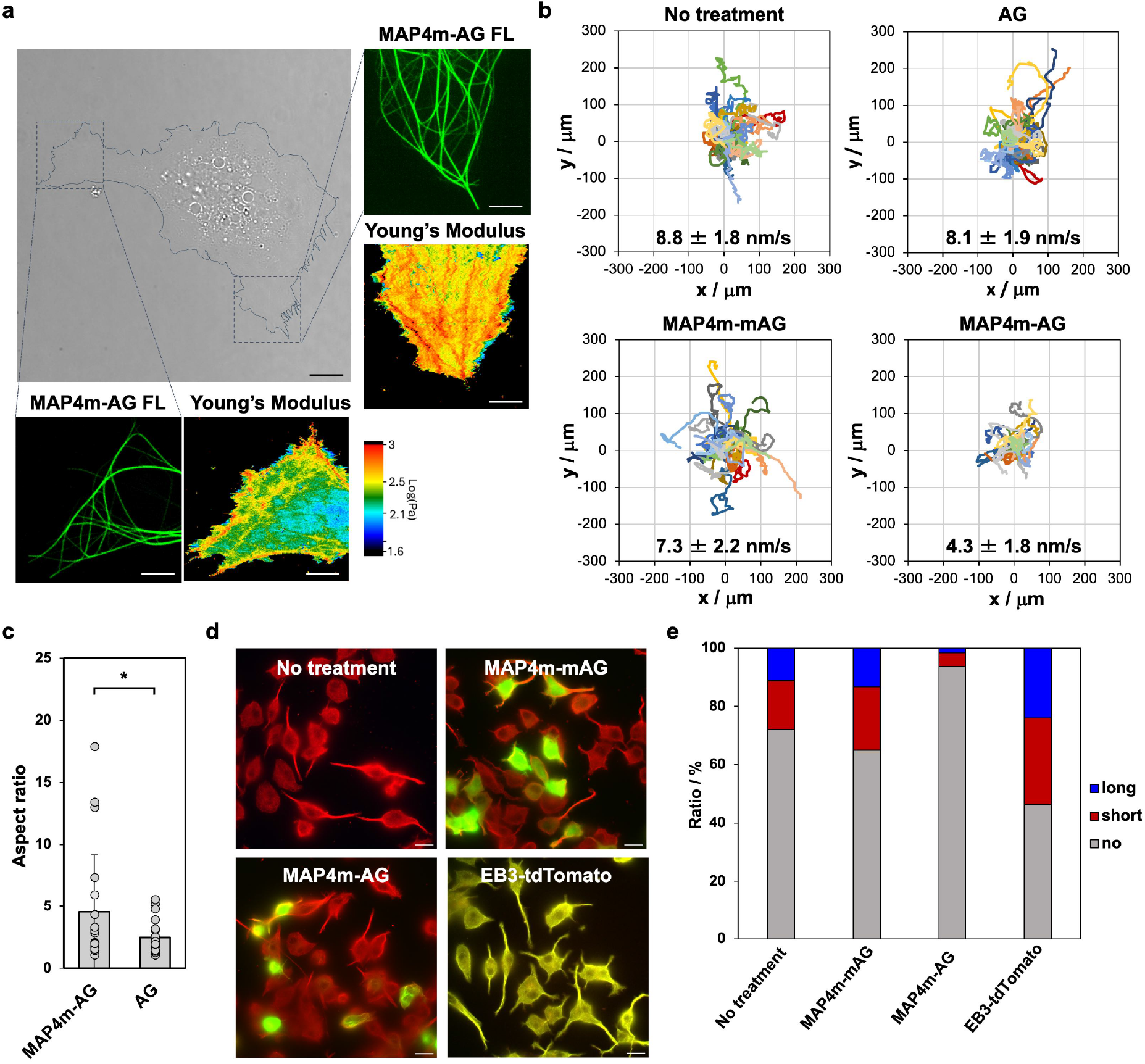
MT-Bundler causes elongation of cell shape, inhibition of cell migration, and reduced neurite/axon formation. (a) Imaging of mechanical properties imaging of cells expressing MAP4m-AG using scanning ion conductance microscopy (SICM). Scale bars: 10 µm for phase-contrast images and 4 µm for confocal and Young’s modulus images. (b) Single-cell tracking of HeLa cells expressing AG, MAP4m-mAG, MAP4m-AG, or not expressing any. Values show mean velocity of each sample. Error values represent the SD (*N* = 30). (c) Aspect ratio of HeLa cells expressing MAP4m-AG or AG (Supplementary Fig. 12). Error bars represent the SD (*N* = 20). \**P* < 0.05, two-tailed Student’s t-test. (d) Fluorescence images of Neuro-2a cells expressing MAP4m-mAG, MAP4m-AG, EB3-tdTomato (positive control), or not expressing any. Neuro-2a cells transfected without or with MAP4m-mAG or MAP4m-AG (green) were immunostained for α-tubulin (red). Neuro-2a cells transfected with EB3-tdTomato were immunostained for α-tubulin (green). Scale bars, 10 μm. (e) The graph shows the neurite length quantified from the fluorescence images. Neurites with lengths of one to two cell bodies were categorized as short neurites, whereas those longer than two cell bodies were counted as long neurites (*N* = 60).

### Temporal control of MT bundling

To transition from correlative observations to causal mechanisms, we engineered inducible variants of MT-Bundler that permit precise spatiotemporal control. We employed both chemically inducible dimerization (CID) and light-inducible dimerization (LID) strategies to modulate the MT bundling state in living cells. For the CID system, we used the rapamycin-induced interaction between FKBP and FRB to physically decouple the bundling components^39,40^. MT-Bundler was split into a cytosolic scaffold (AG-FKBP) and an MT-anchored binding motif (mCerulean3-FRB-MAP4m) (Fig. 5a). Prior to induction, AG-FKBP exhibited a diffuse cytosolic distribution, while the MAP4m fusion localized to the cytoskeleton. Upon rapamycin addition, AG-FKBP was rapidly recruited to MTs, triggering immediate and robust bundling (Fig. 5b, Supplementary Video 4). Quantitative tracking confirmed the rapid accumulation of scaffold fluorescence on MTs, concomitant with the structural reorganization of the network (Fig. 5b, graphs).

**Fig. 5.**
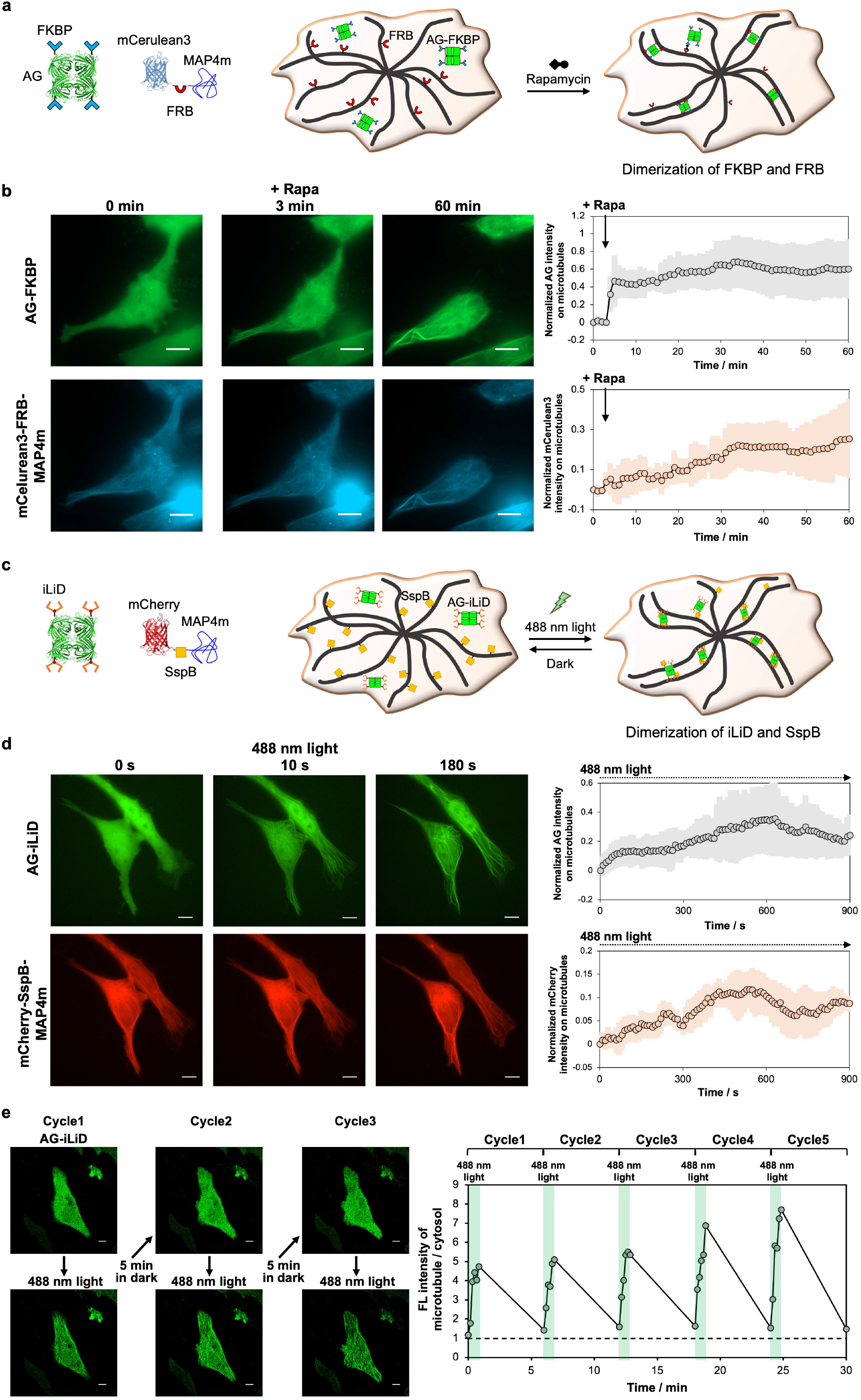
Temporal control of MT bundling. (a) Schematic illustration of MT bundling using FKBP-fused AG and FRB-fused mCeruleaen3-MAP4m by rapamycin treatment. (b) Translocation of AG-FKBP to MTs and induction of bundled MTs by rapamycin treatment (left). Scale bars, 10 μm. Rapa, 100 nM rapamycin. See also Supplementary Video 4. Time-course quantification of the AG-FKBP translocation and normalized mCerulean3 intensity on MTs (right). The colored areas show the SD (*N* = 3). (c) Schematic illustration of light-induced MT bundling using iLiD-fused AG and SspB-fused mCherry-MAP4m. (d) Translocation of AG-iLiD to MTs and induction of the bundling by light irradiation (left). Scale bars, 10 μm. See also Supplementary Video 5. Time-course quantification of the AG-iLiD translocation and normalized mCherry intensity (right). The colored areas show the standard deviation of the mean (*N* = 3). (e) Quantification of reversible dimerization of AG-iLiD and mCherry-SspB-MAP4m.

While CID provides robust induction, it lacks reversibility. To achieve dynamic, reversible control, we developed an optogenetic LID-type MT-Bundler utilizing the iLID/SspB pair, which heterodimerizes upon exposure to 488 nm light and dissociates in the dark^41,42^. We engineered AG-iLID as the scaffold and mCherry-SspB-MAP4m as the anchor (Fig. 5c). Illumination with pulsed 488 nm light drove the rapid recruitment of AG-iLID to MTs, inducing bundle formation comparable to that with the constitutive construct (Fig. 5d, Supplementary Video 5). Crucially, this process proved fully reversible; the cessation of light exposure resulted in dissociation of the scaffold. We confirmed that this bundling–unbundling cycle could be repeated at least five times without loss of responsiveness (Fig. 5e). These genetically encoded actuators thus provide a powerful platform for the acute manipulation of cytoskeletal architecture in living cells.

### Acute bundling triggers acetylation and neurite retraction

Finally, to achieve the temporal control of MT acetylation, we deployed the LID-type MT-Bundler. Following the co-expression of AG-iLID and mCherry-SspB-MAP4m in HeLa cells, we quantified acetylation kinetics in real-time. In contrast to the pre-illumination state, rapid elevation of the level of acetylation was observed within 10 min of pulsed exposure to 488 nm light, reaching a plateau at approximately 30 min (Fig. 6a, b). This fast kinetic profile strongly supports our mechanistic model: Light-induced bundling immediately imposes irregular curvature and lattice defects or expansion, which rapidly facilitate the luminal access of ATAT1 to drive acetylation.

**Fig. 6.**
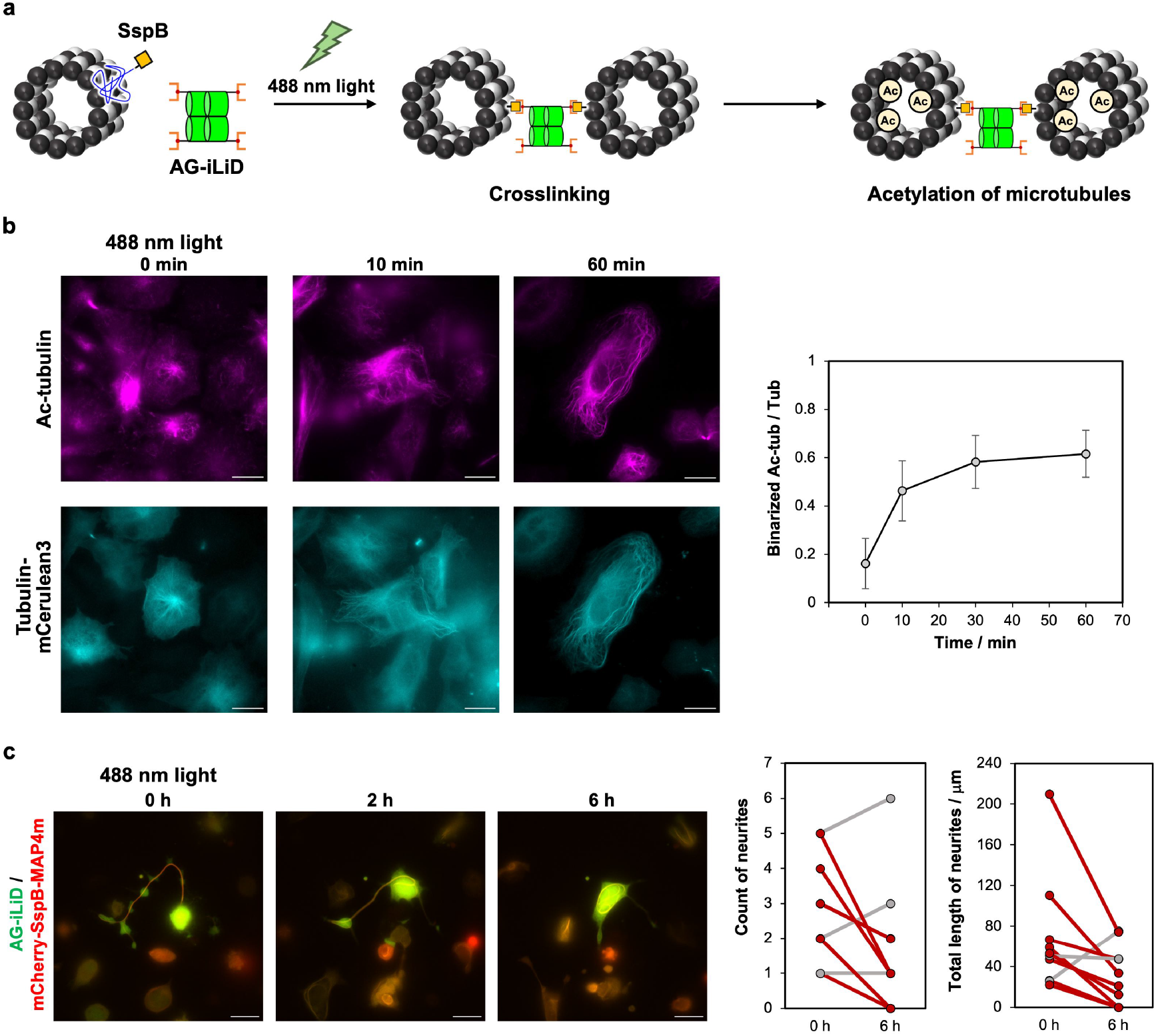
Temporal control of MT acetylation and disruption of neuronal architecture through bundling of MTs. (a) Schematic illustration of light-induced MT acetylation through bundling of MTs using AG-iLiD and mCherry-SspB-MAP4m. (b) Light-induced change of acetylation level of MTs. HeLa cells transfected with AG-iLiD, mCherry-SspB-MAP4m, and tubulin-mCelurean3 were immunostained for Ac-tubulin (left). A 488 nm pulsed laser irradiated the cells every 10 s for 0, 10, or 60 min. Scale bars, 10 μm. Quantification of acetylation level (right). Error bars represent the SD (*N* = 5). (c) Time-lapse images of light-induced inhibition of neurite/axon formation. Scale bars, 10 μm. Graphs show the count of neurites and total neurite length before and after light irradiation every 10 s for 6 h (*N* = 10). Red marks represent a decrease in the count of neurites or total length, while gray marks represent no significant change or an increased in the count of neurites or total length over 6 h.

To determine whether acute bundling is sufficient to disrupt established neuronal architecture, we activated the LID-type MT-Bundler in Neuro-2a cells. Sustained light stimulation (pulsed every 10 s for 6 h) triggered significant neurite retraction (Fig. 6c), effectively phenocopying the morphological defects observed with constitutive MAP4m-AG overexpression. Quantitative analysis confirmed that a large proportion of cells exhibited marked reductions in both neurite number and total length following the acute crosslinking of AG-iLID to MTs. Together, these data establish MT-Bundler as a versatile platform for dissecting the structural logic of the cytoskeleton in living cells.

## DISCUSSION

We developed a genetically encoded MT-bundling tool that enables direct, tunable, and temporally controlled crosslinking of intracellular MT. By decoupling MT bundling from the multifunctional activities of endogenous MAPs, this approach allowed us to examine the intrinsic consequences of MT bundling on MT structure, modification, and cellular behavior. Our results demonstrate that MT bundling alone is sufficient to confer pronounced resistance to depolymerization, even in the absence of MT acetylation. This finding provides direct evidence that physical crosslinking of MTs can stabilize the polymer independently of luminal post-translational modification, a conclusion that has been difficult to establish using endogenous MAPs or pharmacological perturbations. Bundling is therefore not merely correlated with MT stability but represents an autonomous structural mechanism of stabilization.

At the same time, we found that MT bundling promotes the rapid and robust acetylation of α-tubulin. Using optogenetic control, we showed that acetylation follows bundling on a timescale of minutes, placing bundling upstream of ATAT1-mediated modification. Importantly, genetic ablation of ATAT1 abolished acetylation without impairing bundling, further separating these processes. Together, these observations support a model in which MT bundling induces lattice curvature, defects, or expansion that increase luminal accessibility to ATAT1, consistent with recent structural and biochemical studies^23,28,29^. Notably, MT bundling driven by the endogenous MAP Tau conferred ATAT1-independent structural stability, which is consistent with our MT-Bundler data. This correspondence validates our synthetic system as a generalizable model that accurately recapitulates the fundamental functional consequences of physiological MT bundling. These findings clarify a long-standing ambiguity in the field regarding the relationship between MT bundling, stability, and acetylation. While acetylation has often been treated as a proxy for MT stability, our data indicate that acetylation is a downstream consequence of structural reorganization rather than a prerequisite for stabilization. This distinction has important implications for interpreting MT modifications in physiological and pathological contexts.

Functionally, enforced MT bundling profoundly altered cell behavior, including disruptions to mitotic progression, cell migration, and neurite and axon formation. These phenotypes mirror, in exaggerated form, effects attributed to dysregulated MAP activity in both non-neuronal and neuronal systems^11,12,35–38^. In neurons, where precise MT organization underlies axonal polarity and growth, excessive or mislocalized bundling likely interferes with the coordinated actions of endogenous MAPs such as Tau and MAP2. Our results therefore support the idea that not only MT stability but also the spatial and temporal patterning of MT bundling is critical for normal cellular function.

MT-Bundler complements existing strategies for manipulating the MT cytoskeleton. Unlike small-molecule drugs that globally perturb polymerization dynamics, or approaches that directly target modifying enzymes, MT bundling provides a structural mode of intervention that preserves endogenous regulatory pathways. The chemically inducible and optogenetic variants further enable reversible and localized control, opening up the possibility of probing spatially restricted MT architectures, such as perinuclear or Golgi-associated MT networks. Notably, the preferential bundling of dense perinuclear MTs observed here resembles the behavior of endogenous crosslinkers such as MTCL2^8^, suggesting that local MT density may bias bundling efficiency.

Beyond basic cell biology, MT-Bundler may offer a useful framework for exploring disease-relevant MT dysregulation. Reduced MT acetylation and instability are hallmarks of neurodegenerative disorders and certain cancers^43,44^. Because MT bundling stabilizes MTs independently of acetylation, artificial bundling may represent a complementary strategy to restore MT integrity when modifying enzymes are compromised. While such applications remain speculative, the MT-Bundler provides a platform for experimentally testing these ideas.

In summary, our work identifies MT bundling as a primary structural determinant of MT stability and modification and introduces a versatile set of tools to manipulate MT architecture in living cells. By enabling causal dissection of MT organization, MT-Bundler opens up new avenues for understanding how higher-order cytoskeletal structures encode cellular function and dysfunction.

## CONCLUSION

We developed a protein-based tool, MT-Bundler that artificially induces MT bundling in living cells and used it to investigate the role of MT bundling. MT bundling mediated by MT-Bundler enhanced MT structural stability and promoted acetylation. Notably, MT bundling was induced regardless of the acetylation state, and bundling itself directly stabilized MTs. Artificially induced MT bundling exerted diverse effects on cellular behavior, including cell elongation, defective migration, and reduced neurite formation. By engineering chemically inducible (CID-type) and light-inducible (LID-type) MT-Bundlers, we achieved temporal control of MT bundling. Furthermore, the LID-type MT-Bundler enabled light-triggered induction of MT acetylation and neurite contraction. Collectively, our findings suggest that endogenous MT-bundling MAPs may similarly regulate MT structural stability and acetylation. By artificially inducing MT bundling, we were able to clarify, for the first time, aspects of the relationship between MT bundling and acetylation. MT-Bundler provides a versatile tool that should facilitate future studies of MAPs involved in MT bundling as well as proteins regulating MT acetylation, offering a new experimental approach to investigate MT biology.

## Declaration of generative AI and AI-assisted technologies in the writing process

During the preparation of this work, the authors used Gemini 3 Pro to improve language and readability. After using this service, the authors reviewed and edited the content as needed and take full responsibility for the content of the publication.

## ACKNOWLEDGMENTS

We thank N. Matsuura and the technical staff of the Chemical Bio-Life Division, Technical Department, Tottori University, and H. Uga from the University of Tokyo for technical assistance. We thank T. Tomita (The University of Tokyo) for providing Neuro-2a cells. This work was supported by the Japan Science and Technology Agency (JST) FOREST (JPMJFR2034 to H.I., JPMJFR203K to Y.T., and JPMJFR226R to T.S.); the Japan Society for the Promotion of Science (JSPS) KAKENHI (JP23K04931 and JP24H01721 in Grant-in-Aid for Transformative Research Areas “Materials Science of Meso-Hierarchy” to H.I.; JP25KJ1837 to S.W.; 23K27318 to T.C.; and JP23H05480 to Y.T.); AMED-CREST (JP23gm1410012 to Y.T.); JST CREST (JPMJCR24T6 to Y.T.); the Mitsubishi Foundation (to H.I.); the JSPS Overseas Challenge Program for Young Researchers (to S.W.); the World Premier International Research Center Initiative (WPI) Program (MEXT, Japan, to Y.T.); Human Frontier Science Program (HFSP, RGP0028/2022 to Y.T.); and National Institute of Health (NIH, R35GM149329 to T.I.). We are also grateful to Edanz (https://jp.edanz.com/ac) for editing a draft of this manuscript.

## REFERENCES

1. Hirokawa, N. Microtubule organization and dynamics dependent on microtubule-associated proteins. Curr. Opin. Cell Biol. 6, 74–81 (1994).

2. Watanabe, T., Noritake, J. & Kaibuchi, K. Regulation of microtubules in cell migration. Trends Cell Biol. 15, 76–83 (2005).

3. Akhmanova, A. & Steinmetz, M. O. Control of microtubule organization and dynamics: two ends in the limelight. Nat. Rev. Mol. Cell Biol. 16, 711–726 (2015).

4. Brouhard, G. J. & Rice, L. M. Microtubule dynamics: an interplay of biochemistry and mechanics. Nat. Rev. Mol. Cell Biol. 19, 451–463 (2018).

5. Akhmanova, A. & Kapitein, L. C. Mechanisms of microtubule organization in differentiated animal cells. Nat. Rev. Mol. Cell Biol. 23, 541–558 (2022).

6. Glotzer, M. The 3Ms of central spindle assembly: microtubules, motors and MAPs. Nat. Rev. Mol. Cell Biol. 10, 9–20 (2009).

7. Balabanian, L., Berger, C. L. & Hendricks, A. G. Acetylated Microtubules Are Preferentially Bundled Leading to Enhanced Kinesin-1 Motility. Biophys. J. 113, 1551–1560 (2017).

8. Matsuoka, R. et al. MTCL2 promotes asymmetric microtubule organization by crosslinking microtubules on the Golgi membrane. J. Cell Sci. 135, jcs259374 (2022).

9. Halpain, S. & Dehmelt, L. The MAP1 family of microtubule-associated proteins. Genome Biol. 7, 224 (2006).

10. Bodakuntla, S., Jijumon, A. S., Villablanca, C., Gonzalez-Billault, C. & Janke, C. Microtubule-Associated Proteins: Structuring the Cytoskeleton. Trends Cell Biol. 29, 804–819 (2019).

11. Dehmelt, L. & Halpain, S. The MAP2/Tau family of microtubule-associated proteins. Genome Biol. 6, 204 (2004).

12. Kapitein, L. C. & Hoogenraad, C. C. Building the Neuronal Microtubule Cytoskeleton. Neuron 87, 492–506 (2015).

13. Doki, C. et al. Microtubule elongation along actin filaments induced by microtubule-associated protein 4 contributes to the formation of cellular protrusions. J. Biochem. 168, 295–303 (2020).

14. Chapin, S. J. & Chloë Bulinski, J. Non-neuronal 210×103Mr microtubule-associated protein (MAP4) contains a domain homologous to the microtubule-binding domains of neuronal MAP2 and tau. J. Cell Sci. 98, 27–36 (1991).

15. Nguyen, H.-L. et al. Overexpression of full-or partial-length MAP4 stabilizes microtubules and alters cell growth. J. Cell Sci. 110, 281–294 (1997).

16. Mollinari, C. et al. PRC1 is a microtubule binding and bundling protein essential to maintain the mitotic spindle midzone. J. Cell Biol. 157, 1175–1186 (2002).

17. Kader, M. A., Satake, T., Yoshida, M., Hayashi, I. & Suzuki, A. Molecular basis of the microtubule-regulating activity of microtubule crosslinking factor 1. PLoS ONE 12, e0182641 (2017).

18. Satake, T. et al. MTCL1 plays an essential role in maintaining Purkinje neuron axon initial segment. EMBO J. 36, 1227–1242 (2017).

19. Abal, M., Andreu, J. M. & Barasoain, I. Taxanes: Microtubule and Centrosome Targets, and Cell Cycle Dependent Mechanisms of Action. Curr. Cancer Drug Targets 3, 193–203 (2003).

20. Jordan, M. A. & Wilson, L. Microtubules as a target for anticancer drugs. Nat. Rev. Cancer 4, 253–265 (2004).

21. Borowiak, M. et al. Photoswitchable Inhibitors of Microtubule Dynamics Optically Control Mitosis and Cell Death. Cell 162, 403–411 (2015).

22. Steinmetz, M. O. & Prota, A. E. Microtubule-Targeting Agents: Strategies To Hijack the Cytoskeleton. Trends Cell Biol. 28, 776–792 (2018).

23. Janke, C. & Montagnac, G. Causes and Consequences of Microtubule Acetylation. Curr. Biol. 27, R1287–R1292 (2017).

24. Deb Roy, A. et al. Non-catalytic allostery in α-TAT1 by a phospho-switch drives dynamic microtubule acetylation. J. Cell Biol. 221, e202202100 (2022).

25. Janke, C. & Chloë Bulinski, J. Post-translational regulation of the microtubule cytoskeleton: mechanisms and functions. Nat. Rev. Mol. Cell Biol. 12, 773–786 (2011).

26. Janke, C. & Magiera, M. M. The tubulin code and its role in controlling microtubule properties and functions. Nat. Rev. Mol. Cell Biol. 21, 307–326 (2020).

27. Roll-Mecak, A. The Tubulin Code in Microtubule Dynamics and Information Encoding. Dev. Cell 54, 7–20 (2020).

28. Coombes, C. et al. Mechanism of microtubule lumen entry for the α-tubulin acetyltransferase enzyme αTAT1. Proc. Natl. Acad. Sci. U.S.A. 113, E7176–E7184 (2016).

29. Egoldt, C., Velluz, M.-C., Tran, J. & Aumeier, C. Microtubule lattice conformation and integrity regulate α-tubulin acetylation. bioRxiv, (2025). DOI: 10.1101/2025.09.09.675099

30. Inaba, H. et al. Generation of stable microtubule superstructures by binding of peptide-fused tetrameric proteins to inside and outside. Sci. Adv. 8, eabq3817 (2022).

31. Watari, S. et al. Optical Control of Microtubule Accumulation and Dispersion by Tau-Derived Peptide-Fused Photoresponsive Protein. JACS Au 5, 791–801 (2025).

32. Nihongaki, Y., Matsubayashi, H. T. & Inoue, T. A molecular trap inside microtubules probes luminal access by soluble proteins. Nat. Chem. Biol. 17, 888–895 (2021).

33. Liu, G. Y. et al. Precise control of microtubule disassembly in living cells. EMBO J. 41, e110472 (2022).

34. Rheinlaender, J. & Schäffer, T. E. Quantifying and Utilizing Electroosmotic Flow for Mechanical Measurements with the Scanning Ion Conductance Microscope. Anal. Chem. 97, 22541–22547 (2025).

35. Ezratty, E. J., Partridge, M. A. & Gundersen, G. G. Microtubule-induced focal adhesion disassembly is mediated by dynamin and focal adhesion kinase. Nat. Cell Biol. 7, 581–590 (2005).

36. Stramer, B. et al. Clasp-mediated microtubule bundling regulates persistent motility and contact repulsion in Drosophila macrophages in vivo. J. Cell Biol. 189, 681–689 (2010).

37. Conde, C. & Cáceres, A. Microtubule assembly, organization and dynamics in axons and dendrites. Nat. Rev. Neurosci. 10, 319–332 (2009).

38. Hsieh, P.-C. et al. DDA3 stabilizes microtubules and suppresses neurite formation. J. Cell Sci. 125, 3402–3411 (2012).

39. DeRose, R., Miyamoto, T. & Inoue, T. Manipulating signaling at will: chemically-inducible dimerization (CID) techniques resolve problems in cell biology. Pflugers Arch -Eur. J. Physiol. 465, 409–417 (2013).

40. Voß, S., Klewer, L. & Wu, Y.-W. Chemically induced dimerization: reversible and spatiotemporal control of protein function in cells. Curr. Opin. Chem. Biol. 28, 194–201 (2015).

41. Zimmerman, S. P. et al. Tuning the Binding Affinities and Reversion Kinetics of a Light Inducible Dimer Allows Control of Transmembrane Protein Localization. Biochemistry 55, 5264–5271 (2016).

42. Klewer, L. & Wu, Y.-W. Light-Induced Dimerization Approaches to Control Cellular Processes. Chem. Eur. J. 25, 12452–12463 (2019).

43. Li, L. et al. ATAT1 regulates forebrain development and stress-induced tubulin hyperacetylation. Cell. Mol. Life Sci. 76, 3621–3640 (2019).

44. Wattanathamsan, O. et al. Inhibition of histone deacetylase 6 destabilizes ERK phosphorylation and suppresses cancer proliferation via modulation of the tubulin acetylation-GRP78 interaction. J. Biomed. Sci. 30, 4 (2023).

