## Supplementary Information for "A genetically encoded microtubule bundler for causal dissection of microtubule bundling in cells"

### Experimental Section

#### Equipment and materials.

Confocal laser scanning microscopy (CLSM) measurement was carried out using a FluoView FV10i and FV4000 (Olympus). Fluorescence microscopy measurement was carried out using a ECLIPSE Ti and Ti2-E inverted fluorescence microscope (Nikon), an Axioplan2 fluorescence microscope (Carl Zeiss), Stochastic Optical Reconstruction Microscopy was carried out using a ECLIPSE Ti (Nikon). The reagents were purchased from Nacalai Tesque, Inc., Dojindo Laboratories Co. Ltd., Wako Pure Chemical Industries, New England Biolabs, Thermo Fisher Scientific Inc., Cell Signaling Technology, Inc., Proteintech Group, Inc. and Sigma-Aldrich and used without further purification.

#### Construction of plasmids.

The MAP4m-AG, MAP4m-mAG, AG-FKBP, AG-iLiD expression vectors were constructed based on the pCMV expression vector (RIKEN, Japan) and generated by HiFi DNA assembly. Complementary DNAs (cDNAs) encoding fragments of MAP4m, FKBP, iLiD were amplified by standard polymerase chain reaction (PCR). The amplified fragments were cloned into the open circular pCMV vector amplified by inverse PCR using HiFi DNA assembly master mix (New England Biolabs). PCR was performed in the Thermal Cycler GeneAtlas (astec) using Q5 polymerase master mix (New England Biolabs). For the MAP4m construct, a human truncated mutant (amino acids 800–1115) containing proline-rich domains and affinity domains was used. All the plasmids were sequenced by Eurofins Genomics (Tokyo, Japan). The protein sequences of the constructs are described in Supplementary Fig. 1.

#### Cell culture.

HeLa (RIKEN BioResource Research Center, Japan) and Neuro-2a cells (gift from Prof. Taisuke Tomita from the university of Tokyo) were maintained in DMEM (Wako 043-30085) supplemented with 10% FBS (Wako) and 1% Antibiotic-Antimycotic (gibco) at 37 °C under 5% $CO_2$. RPE-1 (ATCC, CRL-4000) cells were maintained in DMEM/Ham's F-12 supplemented with 10% FBS (Wako) and 1% Antibiotic-Antimycotic (gibco) at 37 °C under 5% $CO_2$.

#### Live-cell imaging of MAP4m-AG-bound microtubules.

For binding analysis of MAP4m-AG to microtubules, HeLa cells were seeded at a density of 40000 cells per well into single-well glass base dish (AGC TECHNO GLASS CO., LTD.). The transfection of pCMV-MAP4m-AG was performed with Lipofectamine LTX (Thermo Fisher Scientific Inc.) according to the manufacturer’s protocol. At 20 h post transfection, staining microtubules was performed with Tubulin Tracker Deep Red (Thermo Fisher Scientific Inc.) according to the manufacturer's protocol. Nucleus were stained by Hoechst 33342 for 30 min. For line profiling of fluorescence intensity of Tubulin Tracker Deep Red, lines of interest were manually drawn for each sample to measure fluorescence intensity of Tubulin Tracker Deep Red. After background subtraction by rolling ball collection, the fluorescence intensity of Tubulin Tracker Deep Red was measured by Fiji software. For calculation of microtubule area per cell area to show microtubule bundling, cell surroundings were manually drawn for each cell to calculate cell area. The rolling ball correction was used to remove the cytosol background from the RAW images. The images were processed to generate the binary microtubule filament pattern via making threshold. Then microtubules areas were calculated by Fiji software.

**Stochastic optical reconstruction microscopy (STORM) measurement.**

HeLa cells were seeded at a density of 40000 cells per well into single-well glass base dish. The transfection of pCMV-MAP4m-AG was performed with FuGENE HD (Promega) according to the manufacturer's protocol. At 20 h post transfection, cells were washed with PBS(−) three times and fixed with 4% paraformaldehyde (PFA) in PBS(−) for 10 min at room temperature. Next, cells were washed with PBS(−) three times and permeabilized with 0.1% Triton X-100 in PBS(−) for 10 min at room temperature. After permeabilization, cells were washed with PBS(−) three times and blocked with 3% BSA and 0.2% Triton X-100 in PBS(−) for 1 h at room temperature. After blocking, cells were incubated with primary antibody against β-tubulin (abcam ab6160) diluted in 3% BSA and 0.2% Triton X-100 in PBS(−) at 1:100 for 30 min at room temperature. Cells were then washed with 0.2% BSA and 0.05% Triton X-100 in PBS(−) three times and incubated with diluted anti-rat secondary antibody conjugated with Alexa Fluor 647 (Thermo Fisher Scientific Inc. A21247) in PBS(−) for 30 min at room temperature. Cells were then washed with 0.2% BSA and 0.05% Triton X-100 in PBS(−) three times subsequently cells were washed with PBS(−) three times and conducted super-resolution STORM imaging. Super-resolution STORM imaging was performed using an inverted total internal reflection fluorescence (TIRF) microscope (Nikon Ti, Nikon) equipped with a 100× oil immersion objective lens (CFI Plan Apochromat DM Lambda 100X Oil, Nikon). A 647 nm laser (Obis LX 647, Coherent) was used for excitation. Fluorescence images were captured using an EMCCD camera (iXon 3, Andor Technology). The imaging buffer was prepared according to a standard STORM protocol[1]. It contained cysteamine (MEA; #30070-50G, Sigma-Aldrich) as a reducing agent, along with an oxygen scavenging system consisting of glucose oxidase (Type VII from Aspergillus, #G2133-250KU; Sigma-Aldrich) and catalase (from bovine liver, #035-12903; Wako). Image sequences were acquired with an exposure time of 20 ms for 5,000 frames. From these 5,000 image sequences, reconstruction of super-resolution images was performed using the ThunderSTORM software[2].

**Evaluation of stability of intracellular microtubules bound with MAP4m-AG.**

For evaluating stability of microtubules against nocodazole treatment, HeLa cells were seeded at a density of 40000 cells per well into eight-well glass chamber. The transfection of pCMV-MAP4m-AG, pCMV-AG or pCMV-MAP4m-mAG was performed with FuGENE HD (Promega) according to the manufacturer's protocol. At 20 h post transfection, cells were treated with 1 μM nocodazole for 1 h at 37 °C. After nocodazole treatment, cells were washed with PBS(−) three times and fixed with 4% paraformaldehyde (PFA) in PBS(−) for 10 min at room temperature. Next, cells were washed with PBS(−) three times and permeabilized with 0.1% Triton X-100 in PBS(−) for 10 min at room temperature. After permeabilization, cells were washed with PBS(−) three times and blocked with 2% BSA in PBS(−) for 1 h at room temperature. After blocking, cells were incubated with primary antibody against α-tubulin (abcam ab6160) diluted in 2% BSA in PBS(−) at 1:1000 overnight at 4 °C. Cells were then washed with PBS(−) three times and incubated with diluted anti-rat secondary antibody conjugated with Alexa Fluor 594 (Thermo Fisher Scientific Inc. A-11007) in PBS(−) for 1 h at room temperature. Cells were washed with PBS(−) three times and conducted fluorescence imaging performed on an Eclipse Ti inverted fluorescence microscope (Nikon) equipped with ×100 oil-immersion objective lens (Nikon). For evaluating stability of microtubules by incubation on ice, HeLa cells were seeded at a density of 40000 cells per well into eight-well glass chamber. The transfection of pCMV-MAP4m-AG, pCMV-AG or pCMV-MAP4m-mAG was performed with Lipofectamine LTX (Thermo Fisher Scientific Inc.) according to the manufacturer's protocol. At 20 h post transfection, cells were incubated for 25 min on ice. After incubation on ice, cells were washed with PBS(−) three times and fixed with 4% paraformaldehyde (PFA) in PBS(−) for 10 min at room temperature. Next, cells were washed with PBS(−) three times and permeabilized with 0.1% Triton X-100 in PBS(−) for 10 min at room temperature. After permeabilization, cells were washed with PBS(−) three times and blocked with 2% BSA in PBS(−) for 1 h at room temperature. After blocking, cells were incubated with primary antibody against α-tubulin(abcam ab18251) diluted in 2% BSA in PBS(−) at 1:1000 overnight at 4 °C. Cells were then washed with PBS(−) three times and incubated with diluted anti-rabbit secondary antibody conjugated with Alexa Fluor 594 (Thermo Fisher Scientific Inc. A-11012) in PBS(−) for 1 h at room temperature. Cells were washed with PBS(−) three times and conducted fluorescence imaging performed on an Eclipse Ti2-E inverted fluorescence microscope (Nikon) equipped with ×60 oil-immersion objective lens (Nikon). For calculation of microtubule area per cell area to show microtubule stability, RAW images were processed to generate the binary microtubule filament pattern via making threshold. Then ROIs were manually drawn for each cell to calculate ratio microtubule area per cell area and measured by Fiji software.

**Assessment of acetylation level of intracellular microtubules bound with MAP4m-AG.**

HeLa cells were seeded at a density of 40000 cells per well into eight-well glass chamber. The transfection of pCMV-MAP4m-AG, pCMV-AG or pCMV-MAP4m-mAG was performed with FuGENE HD (Promega) according to the manufacturer's protocol. At 20 h post transfection, cells were washed with PBS(−) three times and fixed with 4% paraformaldehyde (PFA) in PBS(−) for 10 min at room temperature. Next, cells were washed with PBS(−) three times and permeabilized with 0.1% Triton X-100 in PBS(−) for 10 min at room temperature. After permeabilization, cells were washed with PBS(−) three times and blocked with 2% BSA in PBS(−) for 1 h at room temperature. After blocking, cells were incubated with primary antibody against acetylated-tubulin (Cell Signaling Technology, Inc. 5335T) diluted in 2% BSA in PBS(−) at 1:1000 overnight at 4 °C. Cells were then washed with PBS(−) three times and incubated with diluted anti-rabbit secondary antibody conjugated with Alexa Fluor 647 (Thermo Fisher Scientific Inc. A-21244) in PBS(−) for 1 h at room temperature. Cells were washed with PBS(−) three times and conducted fluorescence imaging performed on an Eclipse Ti2-E inverted fluorescence microscope (Nikon) equipped with ×100 oil-immersion objective lens (Nikon). For calculation of binarized Ac-tubulin area per tubulin area, cell surroundings were manually drawn for each cell to calculate cell area. The rolling ball correction was used to remove the cytosol background from the RAW images. The images of Ac-tubulin and tubulin channels were processed to generate the binary microtubule filament pattern via making threshold and measured areas by Fiji software.

**Generation of cell lines by retroviral-mediated integration**

To enable the selection of retrovirus transduced cells using puromycin, we knocked out the puromycin acetyltransferase (PAC) gene, which is expressed in RPE-1 cells. An sgRNA targeting PAC (5'-TGTCGAGCCCGACGCGCGTG-3') was synthesized in vitro and Cas9/sgRNA RNPs were electroporated into cells using the Neon Transfection System (Thermo Fischer Scientific) as described previously[3]. Single cells were subsequently isolated into 96-well plates using cellenONE (Cellenion) according to the manufacturer's instructions. RPE-1 PAC KO Tet3G transactivator cell lines were generated and used as parental cell lines for generating Tet-On cell lines as described previously[4]. RPE-1 Tet-On cell lines were established by retroviral-mediated integration. Each pRetroX-TRE3G-MAP4m-AG plasmid and pCMV-VSV-G (Addgene, 8454) were transfected into HEK GP2-293 cells. The medium was harvested and filtered through a 0.45 μm filter (Merck Millipore, SLHVR33RS). Pre-seeded RPE-1 PAC KO Tet3G cells were infected with the virus-containing medium, supplemented with fresh medium, FBS, and 4 μg/mL polybrene (Nacalai Tesque, 12996-81). Single cells were subsequently isolated into 96-well plates using cellenONE (Cellenion) according to the manufacturer's instructions.

**Generation of RPE-1 ATAT1 KO cells**

RPE-1 ATAT1 KO cells were generated by excising the entire ATAT1 gene using a dual-site CRISPR-Cas9 approach. sgRNAs were designed using Custom Alt-R® CRISPR-Cas9 guide RNA (IDT). Single-guide RNAs (sgRNAs) were designed to target the upstream region of first exon (5'-ACTCTGTCAAGATGCCTTGT-3') and the downstream region of last exon (5'-AATGGGTGCAACGACAGTGA-3') of the *ATAT1* locus. These sgRNAs were synthesized by in vitro transcription of DNA oligonucleotide templates using the HiScribe T7 Transcription Kit (New England Biolabs) and subsequently purified with the RNA Clean & Concentrator kit (ZYMO RESEARCH) as described previously[3]. Purified sgRNAs and Alt-R® S.p. HiFi Cas9 Nuclease V3 (IDT, 1081061) were introduced into wild-type RPE1 cells by electroporation using the Neon Transfection System (Thermo Fischer Scientific) as described previously[3]. After several days of culture, single-cell clones were isolated in 96-well plates using cellenONE (Cellenion) according to the manufacturer's instructions. Successful knockout was validated in these clones by genomic PCR (Supplementary Fig. 9) using the following primers: WT detection Fw (5'-AAACTCACCAGAACGAGGTCC-3'), WT detection Rv (5'-GACTCTTGGCACACAAGCCA-3'), KO detection Fw (5'-TCTCTGGTTTGCTTTGGGCT-3'), and KO detection Rv (5'-CCTGCGCGGAACCCAG-3'). The transfection of pCMV-MAP4m-AG into RPE-1 ATAT1 KO was performed with TransIT-X2 reagent (Mirus, MIR 6003) according to the manufacturer's instructions.

**Western blotting analysis.**

Tet-on type RPE1 cells stably expressing MAP4m-AG or MAP4m-mAG were seeded into 10 cm dish. The cells were treated with 1 μg/mL doxycycline and further incubated for 24 h. The cell lysates were prepared by scraping cells using lysis buffer (RIPA buffer, Wako, 188-02453), mixed with protease inhibitor cocktail (Wako, 165-26021). Cell lysates were centrifuged for 10 min at 15,000 g at 4°C to pellet the cell debris, then the supernatants were mixed with 6× Laemmli buffer, heated at 95 °C for 5 min. The mixture was loaded onto SDS-PAGE gel (15% acrylamide) and electrophoresis was carried out at 20 mA (constant current) in a buffer. After electrophoresis, the gel-bound protein bands were electrotransferred to a poly(vinylidene difluoride) membrane. The membrane was blocked by incubation for 1 h at room temperature in Blocking One (10 mL, Nacalai Tesque). The blocked membrane was then incubated with primary antibody against Ac-tubulin (Sigma-Aldrich T6793) and GAPDH (Proteintech Group, Inc., 60004-1) diluted in 10% Blocking One in 1×Tris Buffered Saline (TBS, 5 mM Tris, 13.8 mM NaCl 0.27 mM KCl, pH 7.4) containing 0.1% (v/v) Tween-20 (TBS-T)) for 1 h at room temperature. The membrane was washed with TBS-T and subsequently incubated with horseradish peroxidase-conjugated anti-mouse IgG (Cosmo Bio Co., Ltd.; diluted 1: 10000 with 10% Blocking One in TBS-T) for anti-AC-tubulin and GAPDH. After washing with TBS-T, blots were detected using a chemiluminescence method (ECL; GE

Healthcare). The band images were obtained using a LuminoGraph I CMOS (ATTO). For quantification of acetylation level of MAP4m-AG expressing cell by western blotting, brightness of the Ac-tubulin-derived bands and GAPDH bands were calculated using Fiji. The acetylation level of each sample was calculated by normalizing by GAPDH-derived bands intensity.

**Long-term fluorescence live-cell imaging**

RPE-1 TetOn MAP4m-AG cell lines were cultured in 24-well SENSOPLATE (#662892; Greiner Bio-One) at 37°C in a 5% $CO_2$ atmosphere. 1.5 h before imaging, cells were incubated with 100 nM SiR-tubulin to visualize the microtubules. Time-lapse imaging was carried out with CQ1 Benchtop High-Content Analysis System equipped with a 40× objective (Yokogawa Electric Corp) in the presence of 1 μg/mL doxycycline. Maximum intensity Z-projections of representative images for each condition were generated using the FIJI distribution of the ImageJ software.

**Experiments using Tet-on type RPE1 cells stably expressing MAP4m-AG.**

For calculation of fluorescence intensity of Ac-tubulin with ATAT1 knockout experiments, ROIs were manually drawn for each cell. Fluorescence intensity of Ac-tubulin per tubulin intensity were measured from the fluorescence images by subtracting the background intensity using Fiji software. For calculation of fluorescence intensity of tubulin with ATAT1 knockout experiments, ROIs were manually drawn for each cell. Fluorescence intensities of tubulin were measured from the fluorescence images by subtracting the background intensity using Fiji software. For calculation of microtubule area per cell area to show microtubule stability with ATAT1 knockout experiments, cell surroundings were manually drawn for each cell. The images were processed to generate the binary microtubule filament pattern via making threshold. Then the ratio of microtubule area per cell area were measured by Fiji software.

**Quantification of cell proliferation by WST assay.**

Tet-on type RPE1 cells stably expressing MAP4m-mAG or MAP4m-AG were seeded at a density of 10000 cells per well into 96-well plate. The cells were treated without or with 0.02, 0.2, 2 μg/mL Doxycycline. The cells were further incubated for 36 h. After washing with PBS, the culture medium was replaced with fresh medium (100 μL), and 10 μL of the Cell Counting Kit-8 reaction solution (Dojindo, Japan) was added to each well. After incubation for 4 h, the absorbance at 450 nm was measured. Cell proliferation (%) was calculated by setting the absorbance without doxycycline treatment as 100%.

**Scanning ion-conductance microscopy measurement.**

The setup of the SICM system, scanning algorithms, and experimental protocol have been described previously[5]. The SICM system was mounted on an inverted microscope (Ti2-E, Nikon). The positions of the sample and the nanopipette were controlled using piezoelectric stages in the XY (PKM2H130-040UL, THK Precision) and Z (PS1H45-012UL, THK Precision) directions, driven by a piezo controller (NCM7302CL). Before the measurements were acquired, the culture medium was replaced with L-15 medium. Glass nanopipettes (aperture inner radius 50 nm) filled with L-15 medium were used as probes. Scanning was performed in hopping mode with the following parameters: hopping amplitude, 1.0 μm; waiting time after lateral movement, 1.0 ms (during this time, a reference current was measured as the average of the direct current through the probe); probe approach and withdrawal speed, 80 and 300 nm/ms, respectively; potential +0.6 V, and set point, 98% of the reference current. We recorded ion currents and the vertical probe position as the probe approach the cell surface. We obtained data for a 20 × 20 $\mu m^2$ area containing 256 × 256 points. After the measurements were performed, optical cell images were acquired using a confocal microscopy (CSU-W1 SoRa, Yokogawa) and CFI Plan Apochromat Lambda D 100X Oil objective lens (Nikon), a complementary metal oxide semiconductor (CMOS) camera (ORCA Fusion BT C15440-20UP, Hamamatsu). The SICM data were analyzed, and stiffness maps were produced using a handmade program developed based on LabVIEW2025 (National Instruments). The approach curve of each point was plotted as the ion current versus the vertical probe position, and the slope between 98% and 99% of the reference current was determined with a line fit.
Young's modulus of the cell surface was determined as described previously[6] :

$$E(s) = A_{eo} \cdot p_{eo}(\frac{s_\infty}{s} - 1)^{-1}$$

$$p_{eo} = \frac{8}{3}\frac{\mu_{eo}\eta}{r_i}V_0$$

$E(s)$ is the Young's modulus, $A_{eo}$ is the geometrical parameter of the pipette, and $p_{eo}$ is the electroosmotic pressure. The electroosmotic mobility was assumed to be $\mu_{eo} \approx 2 \times 10^{-8}$ $m^2\,V^{-1}\,s^{-1}$, and the viscosity of the electrolyte was $\eta \approx 1.0$ Pa·s. The inner radius of the nanopipette was $r_i = 50$ nm. The parameters $s$ and $s_\infty$ represent the measured slopes on the cell surface and the rigid substrate, respectively. In the present experiments, $A_{eo}$ was set to 0.12, corresponding to a pipette cone angle of 4°, and $p_{eo}$ was estimated to 12.8 kPa. These parameters were used to calculate the Young's modulus.

**Calculating cell shapes.**

HeLa cells were seeded at a density of 40000 cells per well into eight-well glass chamber. The transfection of pCMV-MAP4m-AG was performed with FuGENE HD (Promega) according to the manufacturer's protocol. At 20 h post transfection, time-lapse imaging was conducted on an

Eclipse Ti inverted fluorescence microscope (Nikon) equipped with ×40 oil-immersion objective lens (Nikon) at 37 °C, 5% $CO_2$ and more than 90% humidity using a stage-top incubator (Tokai Hit) at 10-min intervals for 15 h. For calculation of aspect ratio of cell shapes, length of long side and short side of cells were measured by Fiji software.

**Tracking cell migration.**

HeLa cells were seeded at a density of 20000 cells per well into 12-well plate. The transfection of pCMV-MAP4m-AG was performed with FuGENE HD (Promega) according to the manufacturer's protocol. At 20 h post transfection, time-lapse imaging was conducted on an Eclipse Ti inverted fluorescence microscope (Nikon) equipped with ×10 oil-immersion objective lens (Nikon) at 37 °C, 5% $CO_2$ and more than 90% humidity using a stage-top incubator (Tokai Hit) at 10-min intervals for 15 h. For tracking cell migrations, the location, velocity and total move length were measured by MTrackJ plugin of Fiji software.

**Evaluation of neurite/axon formation of Neuro-2a cells.**

Neuro-2a cells were seeded at a density of 40000 cells per well into eight-well glass chamber. The transfection of pCMV-MAP4m-AG, pCMV-MAP4m-mAG or pCMV-EB3-tdTomato was performed with Lipofectamine LTX (Thermo Fisher Scientific Inc.) according to the manufacturer's protocol. At 48 h post transfection, cells were washed with PBS(−) three times and fixed with 4% paraformaldehyde (PFA) in PBS(−) for 10 min at room temperature. Next, cells were washed with PBS(−) three times and permeabilized with 0.1% Triton X-100 in PBS(−) for 10 min at room temperature. After permeabilization, cells were washed with PBS(−) three times and blocked with 2% BSA in PBS(−) for 1 h at room temperature. After blocking, cells were incubated with primary antibody against α-tubulin (abcam ab18251) diluted in 2% BSA in PBS(−) at 1:1000 overnight at 4 °C. Cells were then washed with PBS(−) three times and incubated with diluted anti-rabbit secondary antibody conjugated with Alexa Fluor 594 (Thermo Fisher Scientific Inc. A-11012) in PBS(−) for 1 h at room temperature. Cells were washed with PBS(−) three times and conducted fluorescence imaging performed on an Eclipse Ti2-E inverted fluorescence microscope (Nikon) equipped with ×60 oil-immersion objective lens (Nikon). For calculation of neurite/axon length, neurite lengths were measured by Fiji software. Neurites with lengths of 1–2 cell bodies were categorized as short neurites, whereas those longer than two cell bodies were counted as long neurites.

**Induction of FKBP-FRB dimerization for formation of bundling microtubules.**

For induction of the FKBP–FRB interaction, HeLa cells were seeded at a density of 40000 cells per well into eight-well glass chamber. The transfection of pCMV-AG-FKBP and pC1-

mCelurean3-FRB-MAP4m was performed with FuGENE HD (Promega) according to the manufacturer's protocol. At 20 h post transfection, the cells were treated with 100 nM rapamycin. Time-lapse imaging was conducted at 37 °C, 5% CO2 and more than 90% humidity using a stage-top incubator (Tokai Hit) at 30-sec intervals for 1 h. For the AG-FKBP colocalization subsequent bundling microtubule analysis, ROIs were manually drawn for each cell. The ROIs were positioned to include mCelurean3-positive microtubules in the peripheral region. The intensity of AG-FKBP and mCelurean3-FRB-MAP4m were calculated by Fiji software. The average value of the normalized AG-FKBP intensity from three different ROIs was used to determine the kinetics of AG-FKBP translocation. The average value of the normalized mCelurean3-FRB-MAP4m intensity from three different ROIs was used to determine the kinetics of microtubule bundling formation.

**Induction of iLID-SspB dimerization for formation of bundling microtubules.**

For induction of the iLiD-SspB interaction for temporal control of microtubule bundling, HeLa cells were seeded at a density of 40000 cells per well into single-well glass base dish. The transfection of pCMV-AG-iLiD and pN1-mCherry-SspB-MAP4m was performed with FuGENE HD (Promega) according to the manufacturer's protocol. At 20 h post transfection, the cells were irradiated 488 nm pulse laser at 10-sec intervals for 30 min. For induction of the iLiD-SspB interaction for showing reversible dimerization, HeLa cells were irradiated 488 nm pulse laser at 10-sec intervals for 1 min and rest in dark for 5 min. For induction of the iLiD-SspB interaction for inhibition of neurite/axon formation, Neuro-2a cells were irradiated 488 nm pulse laser at 10-sec intervals for 15 h.

For the AG-iLiD colocalization subsequent bundling microtubule analysis, ROIs were manually drawn for each cell. The ROIs were positioned to include mCherry-positive microtubules in the peripheral region. The intensity of AG-iLiD and mCherry-SspB-MAP4m were calculated by Fiji software. The average value of the normalized AG-iLiD intensity from three different ROIs was used to determine the kinetics of AG-iLiD translocation. The average value of the normalized mCherry-SspB-MAP4m intensity from three different ROIs was used to determine the kinetics of microtubule bundling formation.

For promotion of acetylation level by light stimulation, HeLa cells were seeded at a density of 40000 cells per well into single well glass base dish. The transfection of pCMV-AG-iLiD and pN1-mCherry-SspB-MAP4m was performed with FuGENE HD (Promega) according to the manufacturer's protocol. At 20 h post transfection, conducted fluorescence imaging performed on an Eclipse Ti2-E inverted fluorescence microscope (Nikon) equipped with ×100 oil-immersion objective lens (Nikon). During time-lapse imaging, 488 nm pulse laser was irradiated to the cells with intervals 10 sec for 0, 10, 30, 60 min. After light stimulation, cells were fixed with cold methanol immediately for 10 min at 4 °C. Next, cells were washed with PBS(−) three times and blocked with 2% BSA in PBS(−) for 1 h at room temperature. After blocking, cells were incubated with primary antibody against Ac-tubulin (Sigma-Aldrich T6793) diluted in 2% BSA in PBS(−) at 1:1000 overnight at 4 °C. Cells were then washed with PBS(−) three times and incubated with diluted anti-mouse secondary antibody conjugated with Alexa Fluor 647 (Thermo Fisher Scientific Inc. A-21235) in PBS(−) for 1 h at room temperature. Cells were washed with PBS(−) three times and conducted fluorescence imaging performed on an Eclipse Ti2-E inverted fluorescence microscope (Nikon) equipped with ×100 oil-immersion objective lens (Nikon).

For calculation of acetylation level with iLiD-SspB dimerization, cell surroundings were manually drawn for each cell to calculate cell area. The rolling ball correction was used to remove the cytosol background from the RAW images. The images of Ac-tubulin and tubulin channels were processed to generate the binary microtubule filament pattern via making threshold and measured areas by Fiji software. For calculation of neurite/axon property, the number of neurite/axons manually counted of each cell. The total lengths of neurite before and after 488 nm pulse laser stimulation intervals 10 sec for 6 h were measured by Fiji software.

**MAP4m-AG (MAP4m(M800-I1115)-AG)**

MSGSKSTQTVAKTTTAAAVASTGPSSRSPSTLLPKKPTAIKTEGKPAEVKKMTAKSVPADLSRPKSTSTSSMKKTTTLSGTAPAAGVVPSRVKATPMP
SRPSTTPFIDKKPTSAKPSSTTPRLSRLATNTSAPDLKNVRSKVGSTENIKHQPGGGRVQIVSKKVSYSHIQSKCGSKDNIKHVPGGGNVQIQNKKVD
ISKVSSKCGSKANIKHKPGGGDVKIESQKLNFKEKAQAKVGSLDNVGHLPAGGAVKTEGGGSEAPLCPGPPAGEEPAISEAAPEAGAPTSASGLNGHP
TLSGGGDQREAQTLDSQIQETSISGLRSRAQASNMVSVIKPEMKIKLCMRGTVNGHNFVIEGEGKGNPYEGTQILDLNVTEGAPLPFAYDILTTVFQY
GNRAFTKYPADIQDYFKQTFPEGYHWERSMTYEDQGICTATSNISMRGDCFFYDIRFDGVNFPPNGPVMQKKTLKWEPSTEKMYVRDGVLKGDVNMAL
LLEGGGHYRCDFKTTYKAKKDVRLPDYHFVDHRIEILKHDKDYNKVKLYENAVARYSMLPSQAK

**AG-MAP4m (AG-MAP4m)**

MVSVIKPEMKIKLCMRGTVNGHNFVIEGEGKGNPYEGTQILDLNVTEGAPLPFAYDILTTVFQYGNRAFTKYPADIQDYFKQTFPEGYHWERSMTYED
QGICTATSNISMRGDCFFYDIRFDGTNFPPNGPVMQKKTLKWEPSTEKMYVRDGVLKGDVNMALLLEGGGHYRCDFKTTYKAKKDVRLPDAHKVDHRI
EILKHDKDYNKVKLYENAVARYSMLPSQAKSGLRSRAQASNMSGSKSTQTVAKTTTAAAVASTGPSSRSPSTLLPKKPTAIKTEGKPAEVKKMTAKSV
PADLSRPKSTSTSSMKKTTTLSGTAPAAGVVPSRVKATPMPSRPSTTPFIDKKPTSAKPSSTTPRLSRLATNTSAPDLKNVRSKVGSTENIKHQPGGG
RVQIVSKKVSYSHIQSKCGSKDNIKHVPGGGNVQIQNKKVDISKVSSKCGSKANIKHKPGGGDVKIESQKLNFKEKAQAKVGSLDNVGHLPAGGAVKT
EGGGSEAPLCPGPPAGEEPAISEAAPEAGAPTSASGLNGHPTLSGGGDQREAQTLDSQIQETSI

**MAP4m-mAG (MAP4m-mAG)**

MSGSKSTQTVAKTTTAAAVASTGPSSRSPSTLLPKKPTAIKTEGKPAEVKKMTAKSVPADLSRPKSTSTSSMKKTTTLSGTAPAAGVVPSRVKATPMP
SRPSTTPFIDKKPTSAKPSSTTPRLSRLATNTSAPDLKNVRSKVGSTENIKHQPGGGRVQIVSKKVSYSHIQSKCGSKDNIKHVPGGGNVQIQNKKVD
ISKVSSKCGSKANIKHKPGGGDVKIESQKLNFKEKAQAKVGSLDNVGHLPAGGAVKTEGGGSEAPLCPGPPAGEEPAISEAAPEAGAPTSASGLNGHP
TLSGGGDQREAQTLDSQIQETSISGLRSRAQASNMVSVIKPEMKIKLCMRGTVNGHNFVIEGEGKGNPYEGTQILDLNVTEGAPLPFAYDILTTVFQY
GNRAFTKYPADIQDYFKQTFPEGYHWERSMTYEDQGICTATSNISMRGDCFFYDIRFDGTNFPPNGPVMQKKTLKWEPSTEKMYVRDGVLKGDVNMAL
LLEGGGHYRCDFKTTYKAKKDVRLPDAHKVDHRIEILKHDKDYNKVKLYENAVARYSMLPSQAK

**Tubulin-mCherry (TUBB5-mCherry)**

MREIVHIQAGQCGNQIGAKFWEVISDEHGIDPTGTYHGDSDLQLDRISVYYNEATGGKYVPRAILVDLEPGTMDSVRSGPFGQIFRPDNFVFGQSGAG
NNWAKGHYTEGAELVDSVLDVVRKEAESCDCLQGFQLTHSLGGGTGSGMGTLLISKIREEYPDRIMNTFSVVPSPKVSDTVVEPYNATLSVHQLVENT
DETYCIDNEALYDICFRTLKLTTPTYGDLNHLVSATMSGVTTCLRFPGQLNADLRKLAVNMVPFPRLHFFMPGFAPLTSRGSQQYRALTVPELTQQVF
DAKNMMAACDPRHGRYLTVAAVFRGRMSMKEVDEQMLNVQNKNSSYFVEWIPNNVKTAVCDIPPRGLKMAVTFIGNSTAIQELFKRISEQFTAMFRRK
AFLHWYTGEGMDEMEFTEAESNMNDLVSEYQQYQDATAEEEEDFGEEAEEEAGGSGCTMVSKGEEDNMAIIKEFMRFKVHMEGSVNGHEFEIEGEGEG
RPYEGTQTAKLKVTKGGPLPFAWDILSPQFMYGSKAYVKHPADIPDYLKLSFPEGFKWERVMNFEDGGVVTVTQDSSLQDGEFIYKVKLRGTNFPSDG
PVMQKKTMGWEASSERMYPEDGALKGEIKQRLKLKDGGHYDAEVKTTYKAKKPVQLPGAYNVNIKLDITSHNEDYTIVEQYERAEGRHSTGGMDELYK

**AG-FKBP (AG-FKBP)**

MVSVIKPEMKIKLCMRGTVNGHNFVIEGEGKGNPYEGTQILDLNVTEGAPLPFAYDILTTVFQYGNRAFTKYPADIQDYFKQTFPEGYHWERSMTYED
QGICTATSNISMRGDCFFYDIRFDGVNFPPNGPVMQKKTLKWEPSTEKMYVRDGVLKGDVNMALLLEGGGHYRCDFKTTYKAKKDVRLPDYHFVDHRI
EILKHDKDYNKVKLYENAVARYSMLPSQAKGGGSGGGSGGSGGGSGGGSGGVQVETISPGDGRTFPKRGQTCVVHYTGMLEDGKKFDSSRDRNKPFKF
MLGKQEVIRGWEEGVAQMSVGQRAKLTISPDYAYGATGHPGIIPPHATLVFDVELLKLE

**mCerulean3-FRB-MAP4m (mCerulean3-FRB-MAP4m)**

MVSKGEELFTGVVPILVELDGDVNGHKFSVSGEGEGDATYGKLTLKFICTTGKLPVPWPTLVTTLSWGVQCFARYPDHMKQHDFFKSAMPEGYVQERT
IFFKDDGNYKTRAEVKFEGDTLVNRIELKGIDFKEDGNILGHKLEYNAIHGNVYITADKQKNGIKANFGLNCNIEDGSVQLADHYQQNTPIGDGPVLL
PDNHYLSTQSKLSKDPNEKRDHMVLLEFVTAAGITLGMDELYKGASGILWHEMWHEGLEEASRLYFGERNVKGMFEVLEPLHAMMERGPQTLKETSFN
QAYGRDLMEAQEWCRKYMKSGNVKDLLQAWDLYYHVFRRISKSAGGSAGGSAGGTGGPRAQASNSGSKSTQTVAKTTTAAAVASTGPSSRSPSTLLPK
KPTAIKTEGKPAEVKKMTAKSVPADLSRPKSTSTSSMKKTTTLSGTAPAAGVVPSRVKATPMPSRPSTTPFIDKKPTSAKPSSTTPRLSRLATNTSAP
DLKNVRSKVGSTENIKHQPGGGRVQIVSKKVSYSHIQSKCGSKDNIKHVPGGGNVQIQNKKVDISKVSSKCGSKANIKHKPGGGDVKIESQKLNFKEK
AQAKVGSLDNVGHLPAGGAVKTEGGGSEAPLCPGPPAGEEPAISEAAPEAGAPTSASGLNGHPTLSGGGDQREAQTLDSQIQETSI

**AG-iLiD (AG-iLiD)**

MVSVIKPEMKIKLCMRGTVNGHNFVIEGEGKGNPYEGTQILDLNVTEGAPLPFAYDILTTVFQYGNRAFTKYPADIQDYFKQTFPEGYHWERSMTYED
QGICTATSNISMRGDCFFYDIRFDGVNFPPNGPVMQKKTLKWEPSTEKMYVRDGVLKGDVNMALLLEGGGHYRCDFKTTYKAKKDVRLPDYHFVDHRI
EILKHDKDYNKVKLYENAVARYSMLPSQAKSAGGSAGGLATTLERIEKNFVITDPRLPDNPIIFASDSFLQLTEYSREEILGRNCRFLQGPETDRATV
RKIRDAIDNQTEVTVQLINYTKSGKKFWNVFHLQPMRDYKGDVQYFIGVQLDGTERLHGAAEREAVCLIKKTAFQIAEAANDENYF

**mCherry-SspB-MAP4m (mCherry-SspB-MAP4m)**

MVSKGEEDNMAIIKEFMRFKVHMEGSVNGHEFEIEGEGEGRPYEGTQTAKLKVTKGGPLPFAWDILSPQFMYGSKAYVKHPADIPDYLKLSFPEGFKW
ERVMNFEDGGVVTVTQDSSLQDGEFIYKVKLRGTNFPSDGPVMQKKTMGWEASSERMYPEDGALKGEIKQRLKLKDGGHYDAEVKTTYKAKKPVQLPG
AYNVNIKLDITSHNEDYTIVEQYERAEGRHSTGGMDELYKGSGSSSPKRPKLLREYYDWLVDNSFTPYLVVDATYLGVNVPVEYVKDGQIVLNLSASA
TGNLQLTNDFIQFNARFKGVSRELYIPMGAALAIYARENGDGVMFEPEEIYDELNIGSGSEFSGSKSTQTVAKTTTAAAVASTGPSSRSPSTLLPKKP
TAIKTEGKPAEVKKMTAKSVPADLSRPKSTSTSSMKKTTTLSGTAPAAGVVPSRVKATPMPSRPSTTPFIDKKPTSAKPSSTTPRLSRLATNTSAPDL
KNVRSKVGSTENIKHQPGGGRVQIVSKKVSYSHIQSKCGSKDNIKHVPGGGNVQIQNKKVDISKVSSKCGSKANIKHKPGGGDVKIESQKLNFKEKAQ
AKVGSLDNVGHLPAGGAVKTEGGGSEAPLCPGPPAGEEPAISEAAPEAGAPTSASGLNGHPTLSGGGDQREAQTLDSQIQETSI

**mAG-Tau382 (mAG-Tau382)**

MVSVIKPEMKIKLCMRGTVNGHNFVIEGEGKGNPYEGTQILDLNVTEGAPLPFAYDILTTVFQYGNRAFTKYPADIQDYFKQTFPEGYHWERSMTYED
QGICTATSNISMRGDCFFYDIRFDGTNFPPNGPVMQKKTLKWEPSTEKMYVRDGVLKGDVNMALLLEGGGHYRCDFKTTYKAKKDVRLPDAHKVDHRI
EILKHDKDYNKVKLYENAVARYSMLPSQAKSAGGSAGGMAEPRQEFEVMEDHAGTYGLGDRKDQGGYTMHQDQEGDTDAGLKAEEAGIGDTPSLEDEA
AGHVTQARMVSKSKDGTGSDDKKAKGADGKTKIATPRGAAPPGQKGQANATRIPAKTPPAPKTPPSSGEPPKSGDRSGYSSPGSPGTPGSRSRTPSLP
TPPTREPKKVAVVRTPPKSPSSAKSRLQTAPVPMPDLKNVKSKIGSTENLKHQPGGGKVQIINKKLDLSNVQSKCGSKDNIKHVPGGGSVQIVYKPVD
LSKVTSKCGSLGNIHHKPGGGQVEVKSEKLDFKDRVQSKIGSLDNITHVPGGGNKKIETHKLTFRENAKAKTDHGAEIVYKSPVVSGDTSPRHLSNVS
STGSIDMVDSPQLATLADEVSASLAKQGL

**Supplementary Fig. 1.** Amino acid sequences of MAP4m-AG, AG-MAP4m, MAP4m-mAG, Tubulin-mCherry, AG-FKBP, mCelurean-FRB-MAP4m, AG-iLiD, mCherry-SspB-MAP4m and mAG-Tau382.

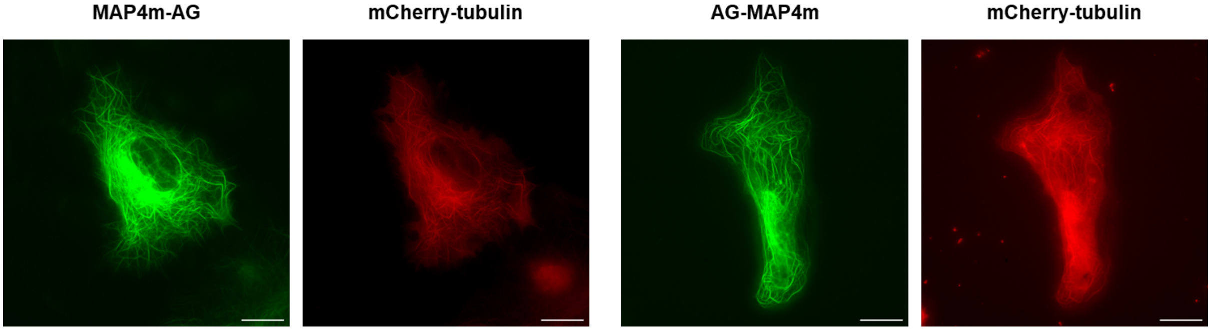


**Supplementary Fig. 2.** Fluorescence images of MAP4m-AG or AG-MAP4m-expressed cells. HeLa cells transfected with MAP4m-AG or AG-MAP4m (green) and mCherry-tubulin (red) for visualizing microtubules. Scale bars, 20 μm.

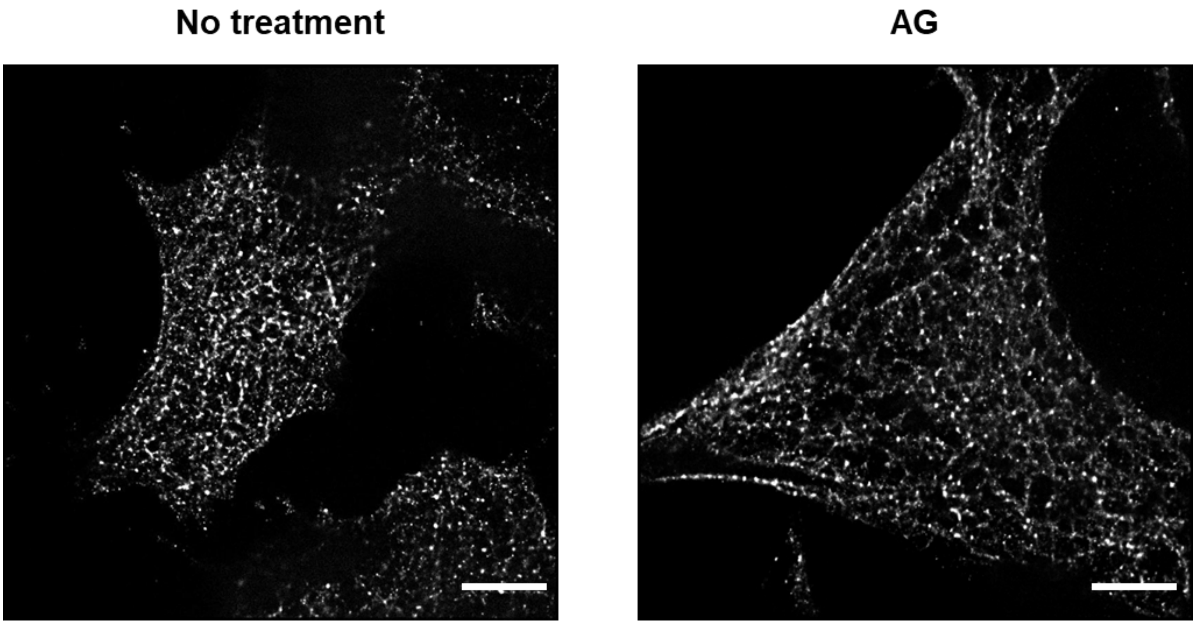


**Supplementary Fig. 3.** Stochastic Optical Reconstruction Microscopy (STORM) images of no treatment or AG-expressed HeLa cells. The samples were stained with β-tubulin antibody. Scale bars, 5 μm.

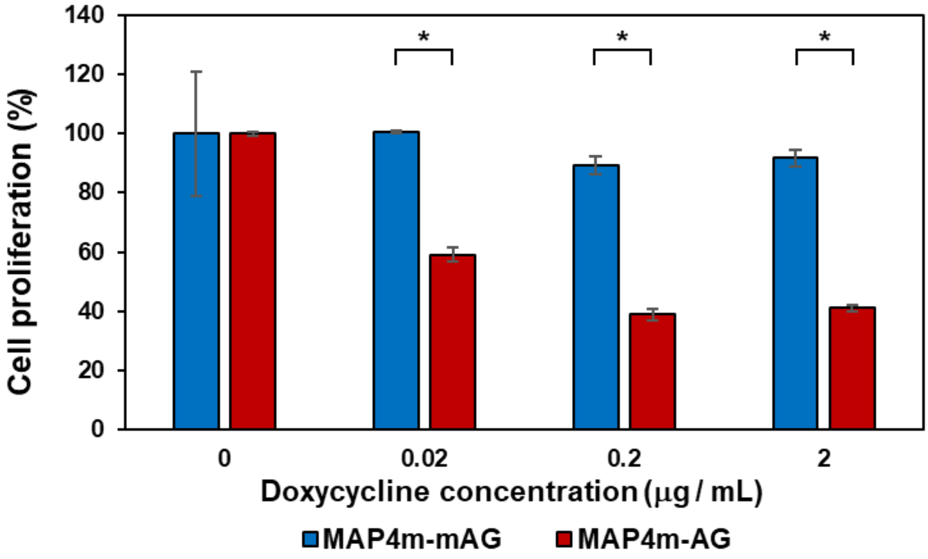


**Supplementary Fig. 4.** Quantification of cell proliferation. Tet-on type RPE1 cells stably expressing MAP4m-mAG or MAP4m-AG were treated without or with 0.02, 0.2, 2 μg/mL Doxycycline. The cells were further incubated for 36 h and cell viability were estimated by WST assay. Error bars represent the SD ($N = 3$). *$P < 0.0005$, two-tailed Student's t-test.

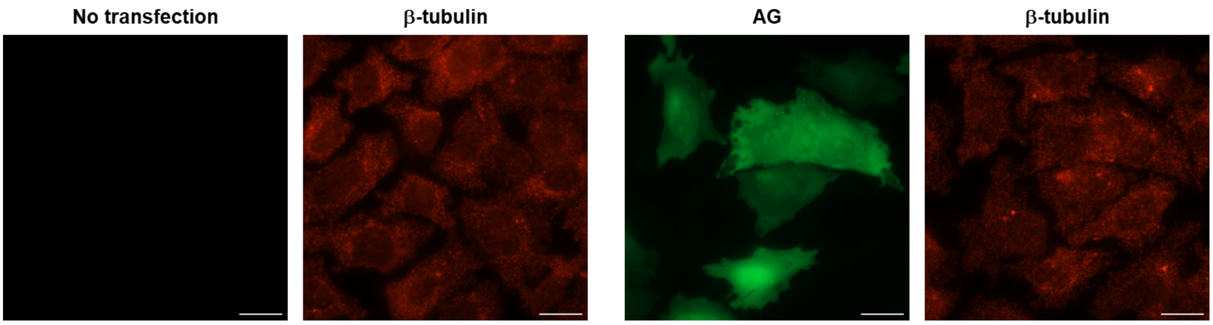


**Supplementary Fig. 5.** Fluorescence images of HeLa cells after 1 μM nocodazole treatment for 2 h. HeLa cells transfected without or with AG (green) were immunostained for β-tubulin (red). Scale bars, 20 μm.

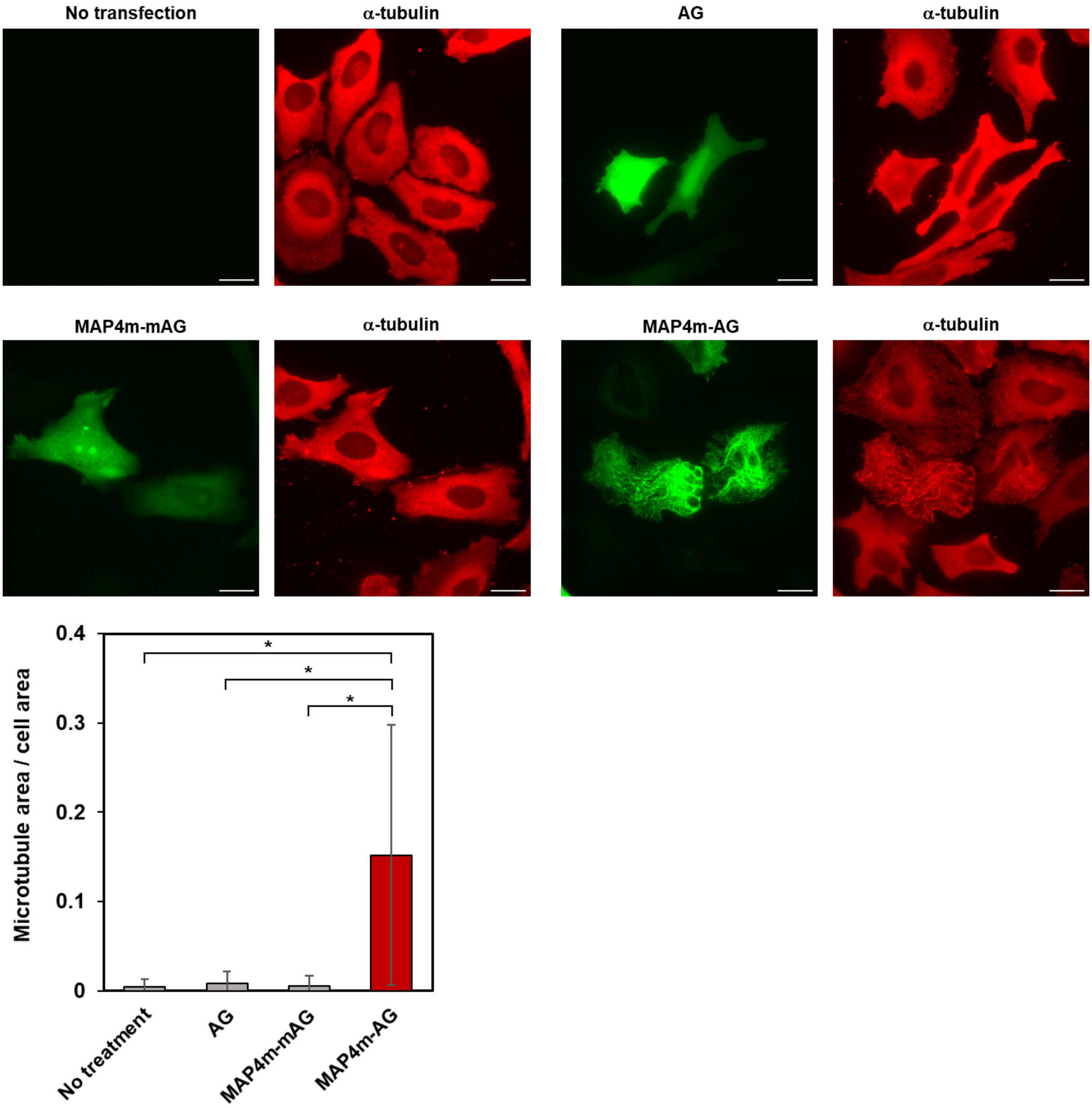


**Supplementary Fig. 6.** Fluorescence images of HeLa cells after incubation on ice for 25 min. HeLa cells transfected without or with AG, MAP4m-mAG, MAP4m-AG (green) were immunostained for α-tubulin (red). Scale bars, 20 μm. A graph shows values of microtubule area per cell area. Error bars represent the SD ($N = 30$). *$P < 0.0001$, two-tailed Student's t-test.

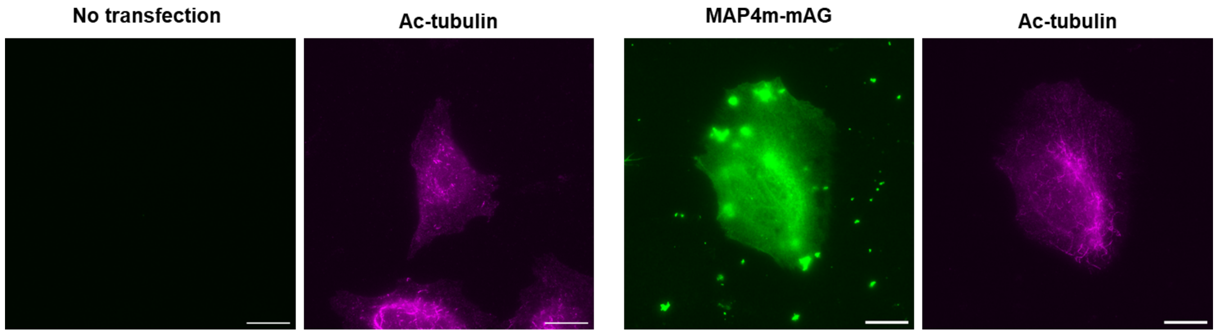


**Supplementary Fig. 7.** Acetylation level of HeLa cells with and without MAP4m-mAG expression. HeLa cells transfected with tubulin-mCherry and MAP4m-mAG (green) were immunostained for acetylated (Ac)-tubulin (magenta). Scale bars, 20 μm.

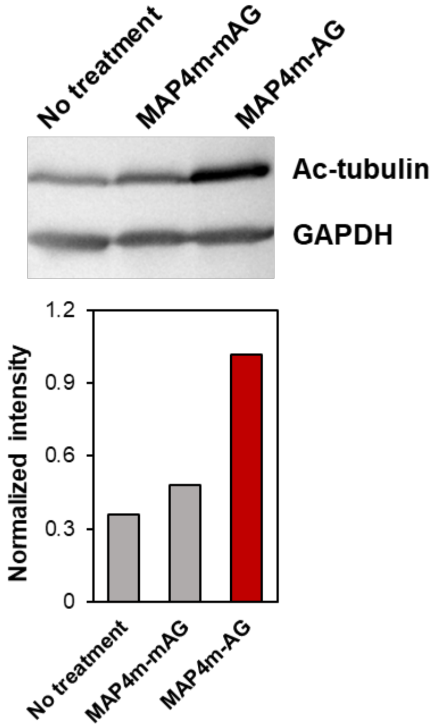


**Supplementary Fig. 8.** Quantification of acetylation level by western blotting. Tet-on type RPE1 cells stably expressing MAP4m-mAG or MAP4m-AG were treated 1 μg/mL Doxycycline. Quantification graph shows intensity of Ac-tubulin normalized by house keeping protein GAPDH.

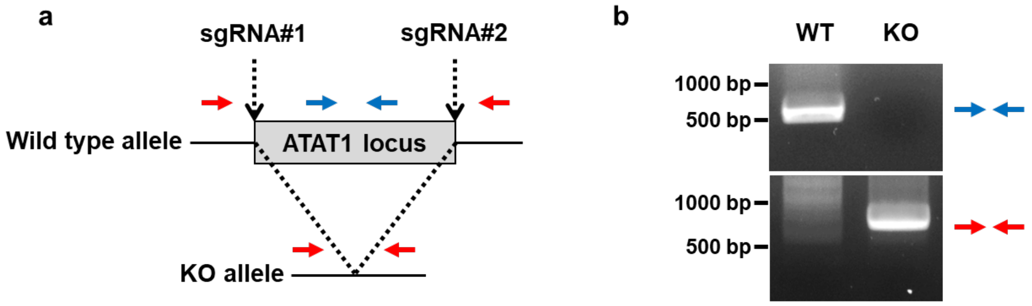


**Supplementary Fig. 9.** (a) Schematic representation of the target positions of sgRNAs and the locations of primers to detect WT (blue) and the deleted region of ATAT1 gene (red), respectively. (b) Genomic PCR for detection of WT and the deleted alleles of ATAT1 gene using the indicated primers, analyzed by agarose gel electrophoresis.

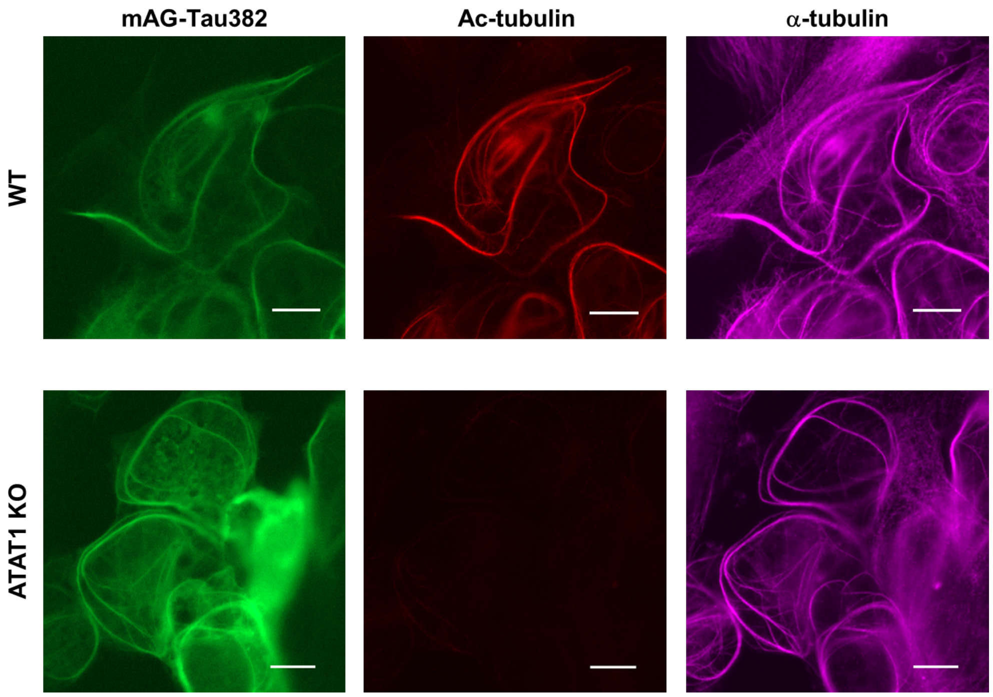


**Supplementary Fig. 10.** Fluorescence images of mAG-Tau382-expressing RPE1 cells with or without ATAT1 knockout. Ac-tubulin and α-tubulin were immunostained. Scale bars, 10 μm.

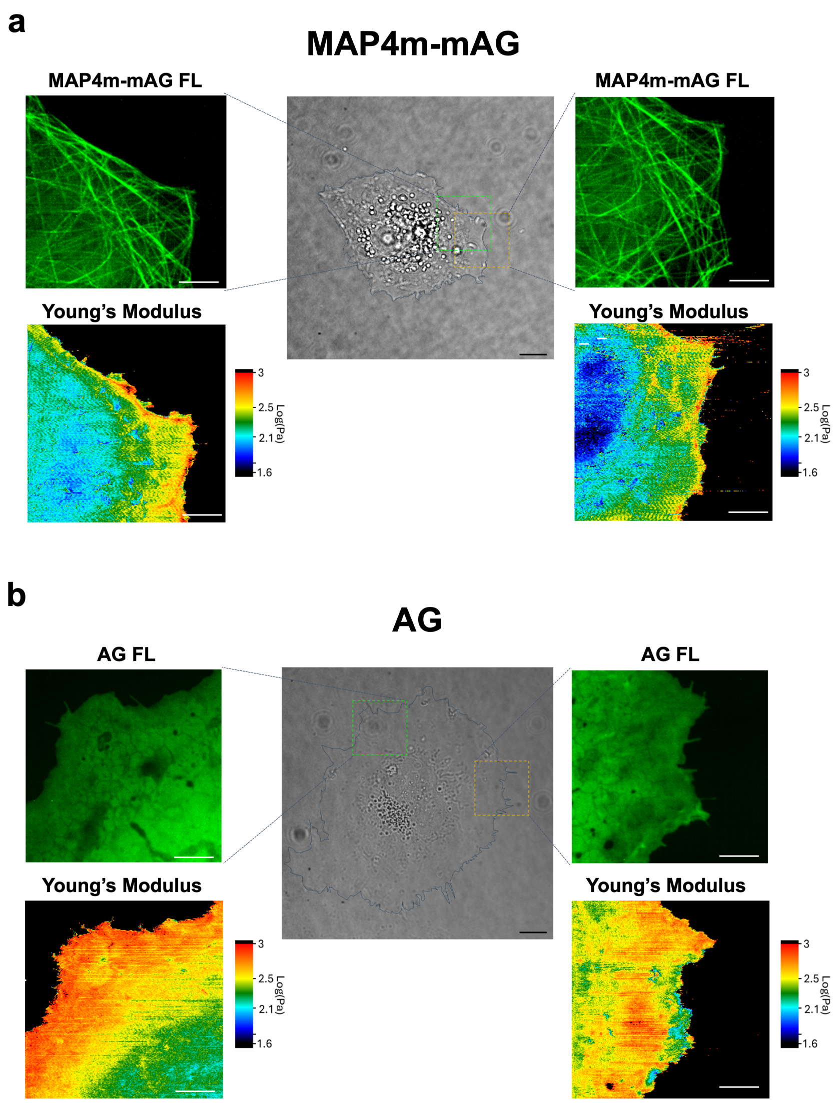


**Supplementary Fig. 11.** Mechanical property imaging of cells expressing MAP4m-mAG (a) or AG (b) using SICM. Scale bars: 10 μm for phase-contrast images and 4 μm for confocal and Young's modulus images.

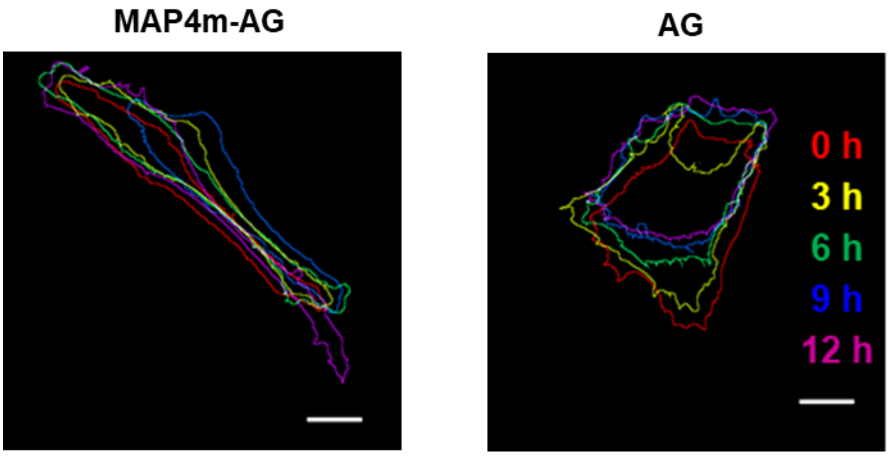


**Supplementary Fig. 12.** (a) Tracking of shapes of HeLa cells expressing MAP4m-AG or AG. Scale bars, 10 μm.

**Supplementary Video 1.** A movie that illustrates the formation of bundling microtubules as MAP4m-AG expressed for 1370 min (Fig. 2a). The intervals are 10 min. The movie is 6000 times faster than the original speed. Scale bar, 50 µm.

**Supplementary Video 2.** A movie that illustrates the tracking of cell migrations with MAP4m-AG expression for 900 min. The intervals are 10 min. The movie is 6000 times faster than the original speed. Scale bar, 100 µm.

**Supplementary Video 3.** A movie that illustrates the change of cell shapes over time by MAP4m-AG expression for 900 min. The intervals are 10 min. The movie is 6000 times faster than the original speed. Scale bar, 20 µm.

**Supplementary Video 4.** A movie that illustrates the translocation of AG-FKBP and subsequent microtubule bundling for 120 min by rapamycin treatment. The intervals are 30 sec. The movie is 300 times faster than the original speed. Scale bar, 10 µm.

**Supplementary Video 5.** A movie that illustrates the translocation of AG-iLiD and subsequent microtubule bundling by 488 nm pulse laser irradiation for 30 min. The intervals are 10 sec. The movie is 100 times faster than the original speed. Scale bar, 10 µm.

**Reference**


1. Dempsey, G. T., Vaughan, J. C., Chen, K. H., Bates, M. & Zhuang, X. Evaluation of fluorophores for optimal performance in localization-based super-resolution imaging. *Nat. Methods* **8**, 1027–1036 (2011).
2. Ovesný, M., Křížek, P., Borkovec, J., Švindrych, Z. & Hagen, G. M. ThunderSTORM: a comprehensive ImageJ plug-in for PALM and STORM data analysis and super-resolution imaging. *Bioinform.* **30**, 2389–2390 (2014).
3. Komori, T. *et al.* A CRISPR-del-based pipeline for complete gene knockout in human diploid cells. *J. Cell Sci.* **136**, jcs260000 (2023).
4. Vlijm, R. *et al.* STED nanoscopy of the centrosome linker reveals a CEP68-organized, periodic rootletin network anchored to a C-Nap1 ring at centrioles. *Proc. Natl. Acad. Sci. U.S.A.* **115**, E2246–E2253 (2018).
5. Takahashi, Y. *et al.* Nanopipette Fabrication Guidelines for SICM Nanoscale Imaging. *Anal. Chem.* **95**, 12664–12672 (2023).

6. Rheinlaender, J. & Schäffer, T. E. Quantifying and Utilizing Electroosmotic Flow for Mechanical Measurements with the Scanning Ion Conductance Microscope. *Anal. Chem.* **97**, 22541–22547 (2025).
